# High-throughput single cell -omics using semi-permeable capsules

**DOI:** 10.1101/2025.03.14.642805

**Authors:** Denis Baronas, Justina Zvirblyte, Simonas Norvaisis, Greta Leonaviciene, Karolis Goda, Vincenta Mikulenaite, Vytautas Kaseta, Karolis Sablauskas, Laimonas Griskevicius, Simonas Juzenas, Linas Mazutis

## Abstract

Biological systems are inherently complex and heterogeneous. Deciphering this complexity increasingly relies on high-throughput analytical methods and tools that efficiently probe the cellular phenotype and genotype. While recent advancements have enabled various single-cell -omics assays, their broader applications are inherently limited by the challenge of efficiently conducting multi-step biochemical assays while retaining various biological analytes. Extending on our previous work (*1*) here we present a versatile technology based on semi-permeable capsules (SPCs), tailored for a variety of high-throughput nucleic acid assays, including digital PCR, genome sequencing, single-cell RNA-sequencing (scRNA-Seq) and FACS-based isolation of individual transcriptomes based on nucleic acid marker of interest. Being biocompatible, the SPCs support single-cell cultivation and clonal expansion over long periods of time – a fundamental limitation of droplet microfluidics systems. Using SPCs we perform scRNA-Seq on white blood cells from patients with hematopoietic disorders and demonstrate that capsule-based sequencing approach (CapSeq) offers superior transcript capture, even for the most challenging cell types. By applying CapSeq on acute myeloid leukemia (AML) samples, we uncover notable changes in transcriptomes of mature granulocytes and monocytes associated with blast and progenitor cell phenotypes. Accurate representation of the entirety of the cellular heterogeneity of clinical samples, driving new insights into the malfunctioning of the innate immune system, and ability to clonally expand individual cells over long periods of time, positions SPC technology as customizable, highly sensitive and broadly applicable tool for easy-to-use, scalable single-cell -omics applications.

## Introduction

Current understanding of the fundamental principles of life increasingly relies on analytical methods and tools that can offer inexpensive and highly accurate single-cell and single-molecule analysis. A rich variety of platforms and techniques developed to-date provide a means to dissect the biological functions of genome (*2-4*), transcriptome (*5-10*), and epigenome (*11-14*) in single cells, and have paved the way for a broad range of applications and biological discoveries (*15-20*). However, existing methods and tools for single-cell -omics research suffer from key inherent constrains. High-throughput methods relying on droplet microfluidics (*5-7*) are largely restricted to one-pot reactions where cells and analytical reagents must be simultaneously loaded during generation of water-in-oil droplets and cannot be replaced downstream the process. On the other hand, microtiter plates offer assay flexibility, yet are cumbersome to use in single-cell biology research, require large reaction volumes, and offer limited throughput. An ideal approach for profiling large numbers of single cells should combine the advantages of both systems – scalability and throughput of droplet microfluidics and versatility of microtiter plate-based platforms.

We recently described the method for generating semi-permeable capsules (SPC) applicable to a high-throughput single-cell RNA cytometry with a single-molecule resolution (*1*). The SPCs, similarly to porous hydrogel beads (*21*), simplify the execution of multi-step reactions through bulk processing, since the required assay reagents can be loaded or removed by simple buffer exchange. However, in contrast to hydrogel beads, where encapsulated cells or their genetic material are entangled within a hydrogel mesh, the SPCs are composite compartments with a liquid core that is wrapped in a thin shell with a tunable permeability. As a result, encapsulated cells and biomolecules that reside in the liquid phase do not experience steric hindrance and obstruction as do biomolecules embedded in a gel phase (*22, 23*). While single genomes, due to their massive size, can be efficiently retained in hydrogels through different enzymatic steps (*24, 25*), capturing smaller entities, such as single cell transcriptomes, is a much more challenging task. Current approaches mainly rely on hybridization of mRNA molecules to poly(dT) primers crafted on a hydrogel mesh (*26, 27*), which yet again negatively impacts transcript capture efficiency and accessibility to nucleic acid processing enzymes.

In this work, we demonstrate a broad utility of SPCs on a variety of nucleic acid assays including digital PCR, single-cell RT-PCR, genome sequencing, single-cell RNA sequencing (scRNA-Seq) by split-and-pool barcoding of mRNA, FACS-based isolation of single transcriptomes based on the expression of RNA marker of interest, and single cell clonal expansion. Once encapsulated in SPCs, tens of thousands of single cells can be separately cultivated and expanded for days and weeks, or alternatively, lysed and processed through a series of biochemical reactions to probe their genotype and phenotype. Taking advantage of these unique properties of SPCs we implemented scRNA-Seq approach (CapSeq) that enables accurate and comprehensive capture of diverse cell types that is challenging to achieve with other state-of-the-art platforms. Single-cell transcriptional profiling of circulating cells of acute myeloid leukemia patients uncovered distinct leukemic cell phenotypes that were shared across the patients and correlated with clinical risk classifications. Furthermore, we identified functionally compromised granulocyte phenotypes expressing the gene markers typically associated with blast and progenitor cells. Next, to accentuate the versatility of SPCs, we combined CapSeq with single-cell RNA cytometry (*1*) and FACS to enrich cells expressing a long non-coding RNA *MIR181A1HG* that was upregulated in AML patients. The accompanying work from Klein and Srivastan groups reports SPCs of different composition and extends their utility beyond scRNA-Seq applications. Taken together, the extensive breadth of applications enabled by the semi-permeable capsules, positioning this technology as the next-generation tool for flexible, user-friendly and scalable single-cell based assays.

## Results

### Physical and biochemical properties of semi-permeable capsules

Semi-permeable capsules (SPCs) are microscopic liquid vessels enveloped by a thin and porous shell (**Figure 1**). We generate SPCs using droplet microfluidics in combination with an aqueous two-phase system (ATPS) composed of dextran and methacryloyl-modified gelatin (GelMA)(*1*). The liquid mixture comprising these two biopolymers exhibits liquid-liquid phase separation phenomenon under a broad range of concentrations (Figure S1). Upon emulsification, the two biopolymers rapidly phase-separate into a concentric core-shell topology, with the polypeptide shell completely enclosing the liquid core (**Figure 1B** and **1C**). To solidify the outer shell, we cool the droplets, remove the carrier oil, and then cross-link the methacryloyl groups by photopolymerization thereby rendering the shell thermo-resistant (**Figure 1D**, Methods). By adjusting the flow rates of the microfluidics device, we obtain SPCs in a size range from 35 to 260 µm, without compromising their concentric structure (Figure S2). Concurrently, by altering the concentration of GelMA we can precisely tune the shell thickness (Figure S2C). Hence, the size of SPCs and shell thickness can be adjusted to meet different experimental needs (Figure S2B and S2C). In a typical scenario we generate the SPCs of 70 ± 2 µm in size and with the shell thickness of 2.2 µm.

**Figure 1.**
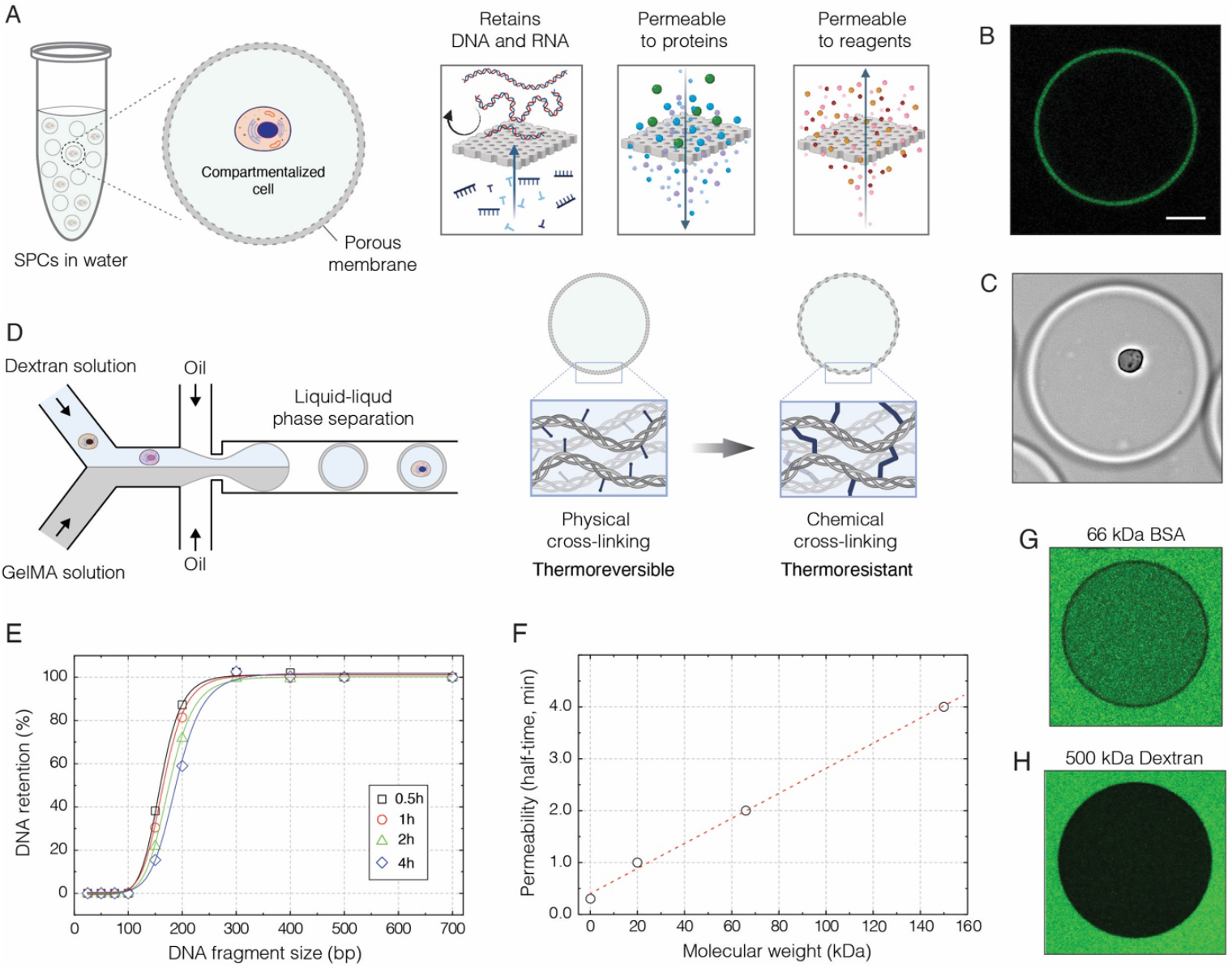
The generation and biochemical properties of semi-permeable capsules (SPCs). **A**) Conceptually, the SPCs represent liquid droplets enveloped by a thin semi-permeable shell, which is reminiscent of a dialysis membrane. This membrane is permeable to assay reagents, oligonucleotides, and enzymes, yet it efficiently retains longer nucleic acid molecules inside the capsule. **B**)The confocal image of a SPC prepared with fluorescently labelled gelatin, which forms a thin outer layer surrounding the liquid core. Scale bar 50 µm. **C**)The bright field image of a SPC suspended in phosphate buffered saline (1x PBS) and carrying a single cell. Scale bar 50 µm. **D**)The generation of SPCs relies on droplet microfluidics and aqueous two-phase system (ATPS), which in this work comprises high molecular weight dextran (500 kDa) and methacryloyl-modified gelatin (GelMA). Upon liquid-liquid phase separation, the GelMA forms a shell, which is then solidified by cooling (physical-crosslinking) followed by photopolymerization (chemical-crosslinking). **E**)Selective retention of nucleic acid fragments by SPCs suspended in 1x PBS, over the course of 4 hours at 22 ºC. **F**)The permeability of SPCs to proteins and biopolymers at 22 ºC. **G**)Fluorescence image of a randomly selected capsule after 3 min incubation with 0.1 µM FITC-BSA (MW 66 kDa) at 22 ºC. **H**)Fluorescence image of a randomly selected capsule after 60 min incubation with 0.2 µM FITC-Dextran (MW 500 kDa) at 22 ºC.

One of the notable characteristics of SPCs is their resilience. The capsules withstand routine laboratory operations, including pipetting, centrifugation, multiple freezing-thawing cycles, and high temperatures (*e*.*g*., 98 ºC). The SPCs remain intact in a variety of organic and inorganic solvents (Figure S3A), yet they can be easily decomposed by protease (Figure S3B). The small amount of dextran (approx. 3% [v/v]) present in the core of SPC, prevents the shell from buckling or collapsing under compression or physical strain, yet, if required, the dextran can be cleared by treating SPCs with dextranase enzyme (Figure S4A).

Another favorable feature of SPCs is their ability to fully retain DNA fragments longer than 300 bp. (**Figure 1E**), while simultaneously allowing assay reagents and proteins to diffuse through (**Figure 1F**). Given the molecular weight of commonly used enzymes such as M-MulV-type reverse transcriptase (71 kDa) and Taq DNA polymerase (97 kDa) these proteins should saturate the core of SPCs within a few minutes (**Figure 1F**). High molecular weight biopolymers such as 500 kDa dextran (Stokes radius of 14.7 nm) cannot permeate SPCs (**Figure 1H**) indicating that the pore size of the shell is below 15 nm. The proteinaceous nature of the shell makes SPCs both biocompatible and biodegradable, while high stability and transparent nature of SPCs makes them amenable to a high-throughput sorting using conventional FACS instruments (Figure S4B), thereby providing a straightforward approach to enrich encapsulated cells for a desirable phenotypic or genotypic trait.

### Semi-permeable capsules provide a versatile tool for complex biochemical and cell-based assays

Given the selective retention of nucleic acids (>300 bp) and permeability to proteins (<200 kDa), SPCs can benefit a variety of biochemical and cell-based assays, some of which are presented in **Figure 2**. In each scenario, cells, microorganisms, nucleic acids, or other biological species are isolated in SPCs at a dilution such that, on average, one capsule carries no more than a single entity. Once sample of interest is encapsulated, the SPCs can be pooled and processed in parallel through a series of biochemical reactions aiming to probe the encapsulated cells and biomolecules (**Figure 2A**). Following this basic principle, we demonstrate a variety of nucleic acid assays such as single DNA molecule amplification and digital quantification (**Figure 2B**), single-cell RT-PCR (**Figure 2C**), single genome amplification (**Figure 2D**), and whole genome sequencing (**Figure 2E**). These non-exhaustive examples illustrate how SPCs can be leveraged to conduct complex nucleic acid assays in regular laboratory tubes while maintaining single-cell, or single molecule, resolution.

**Figure 2.**
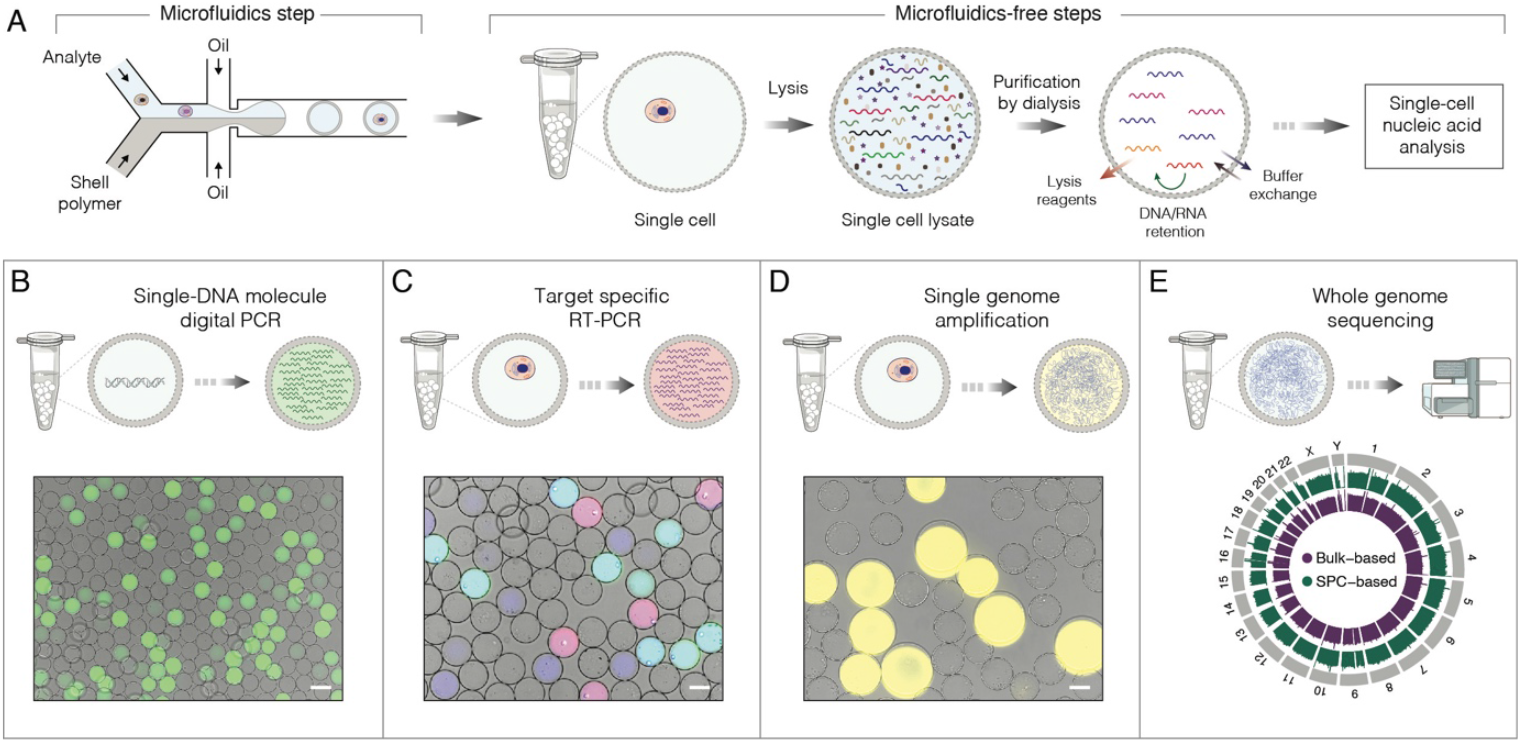
Selected examples of nucleic acid applications using SPCs. **A**) The general strategy for conducting nucleic acid assays on single cells, or single DNA molecules, encapsulated in SPCs. At first, analytes of interest are isolated in SPCs using high-throughput droplet microfluidics followed by subsequent steps that are microfluidics-free and are conducted in a regular laboratory tube by processing all SPCs in bulk. **B**) Single DNA molecule amplification and analysis by digital PCR (staining dye: SYBR Green I). **C**) Multiplex single-cell RT-PCR targeting transcripts encoding CD45, yes-associated protein 1 and β-actin. The SPCs carrying cells expressing both CD45 and ACTB appear as cyan, YAP and ACTB as pink, and ACTB alone as blue. **D**) Single genome amplification of human lymphoblast cells (K-562) by phi29 DNA polymerase and staining with SYTO dye (yellow). The SPCs containing single amplified genomes appear larger due to swelling. **E**) The circular map of a whole genome sequencing of SPC-encapsulated lymphoblast cells. Scale bars, 50 µm

Besides nucleic acid analysis, the SPC technology supports cell growth over a long period of time thereby overcoming a major limitation of droplet microfluidics (**Figure 3A**). For example, in **Figure 3B** we demonstrate a single cell derived spheroid formation in capsule over a period of 11 days, although cultivation for several weeks is also feasible. Similarly, microorganisms such as bacteria and yeast can be effectively expanded into isogenic microcolonies and analyzed further (Figure S4C and S4D). Interestingly, as the dividing cells fill the entire volume of the capsule and continue expanding, the elastic nature of the shell sustains the integrity of SPC allowing it to swell more than 2-times in size, without bursting (**Figure 3C**). A brief exposure to collagenase enzyme, however, rapidly disintegrates the proteinaceous shell thereby enabling fast release of the encapsulated cells without damaging them (**Figure 3C** and Figure S4E, Supplementary Videos 1 and 2). Of note, releasing the inner content of SPCs under mild and physiologically relevant conditions is also important for efficient recovery of encapsulated nucleic acids and other delicate biomolecules. Overall, we show that semi-permeable capsules provide a versatile and universal platform for conducting a broad range of biological and nucleic acid-based assays that are built on multi-step reactions and workflows.

**Figure 3.**
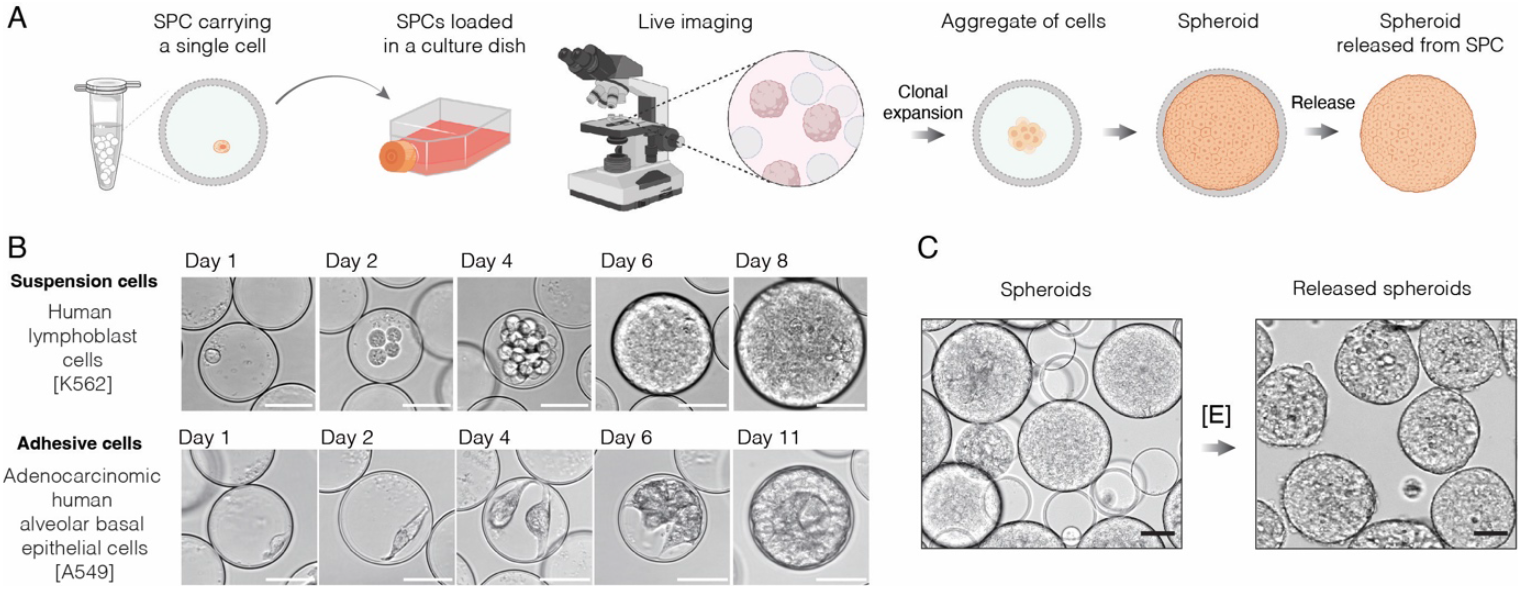
The use of SPCs for live cell cultivation and expansion. **A**) The schematics of conducting 3D cell cultivation and clonal expansion with SPCs. The single cells isolated in SPCs are placed in a suitable growth medium to allow clonal cell expansion and spheroid formation. **B**) The cell culture in SPCs using suspension cells (top row) and adhesive cells (bottom row) over the course of 8 and 11 days, respectively. **C**) The spheroids enclosed in SPCs are released upon addition of collagenase enzyme [E]. Scale bars, 50 µm.

### Capsule-based RNA-Sequencing (CapSeq) for high-throughput single-cell transcriptomics

Considering the broad utility of SPCs, as one of the applications we sought to derive a method for single-cell transcriptomics that we named CapSeq (Capsule-based RNA-Sequencing). The overview of CapSeq is outlined in **Figure 4A**. At first, we isolate single cells in SPCs and following their lysis and several washing steps we obtain SPCs with purified individual transcriptomes free of intracellular inhibitors (e.g., nucleases). We then distribute SPCs in a 96-well plate (∼1000 SPCs per well) and perform cDNA synthesis with poly(dT) primers having a 1^st^ set of barcodes (R1) and unique molecular identifier (UMI). After completing RT step, we pool the SPCs and redistribute them into 2^nd^ microtiter plate and conduct a ligation reaction with a 2^nd^ set of barcodes (R2). We repeat a split-and-pool procedure one more time to tag cDNAs with a 3^rd^ barcode (R3). After completing 3-rounds of barcoding, the SPCs are pooled and redistributed into sub-libraries at a desirable cell count, and sequencing barcodes (R4) are incorporated by indexing PCR. As a result, we can obtain over 10 million barcode combinations (R1-R2-R3-R4) enabling us to barcode 300k of single cells in one run, with 1% multiplet rate. After sequencing, the composition of the cell transcriptome is determined by combining reads with the same barcode combination (Methods).

**Figure 4.**
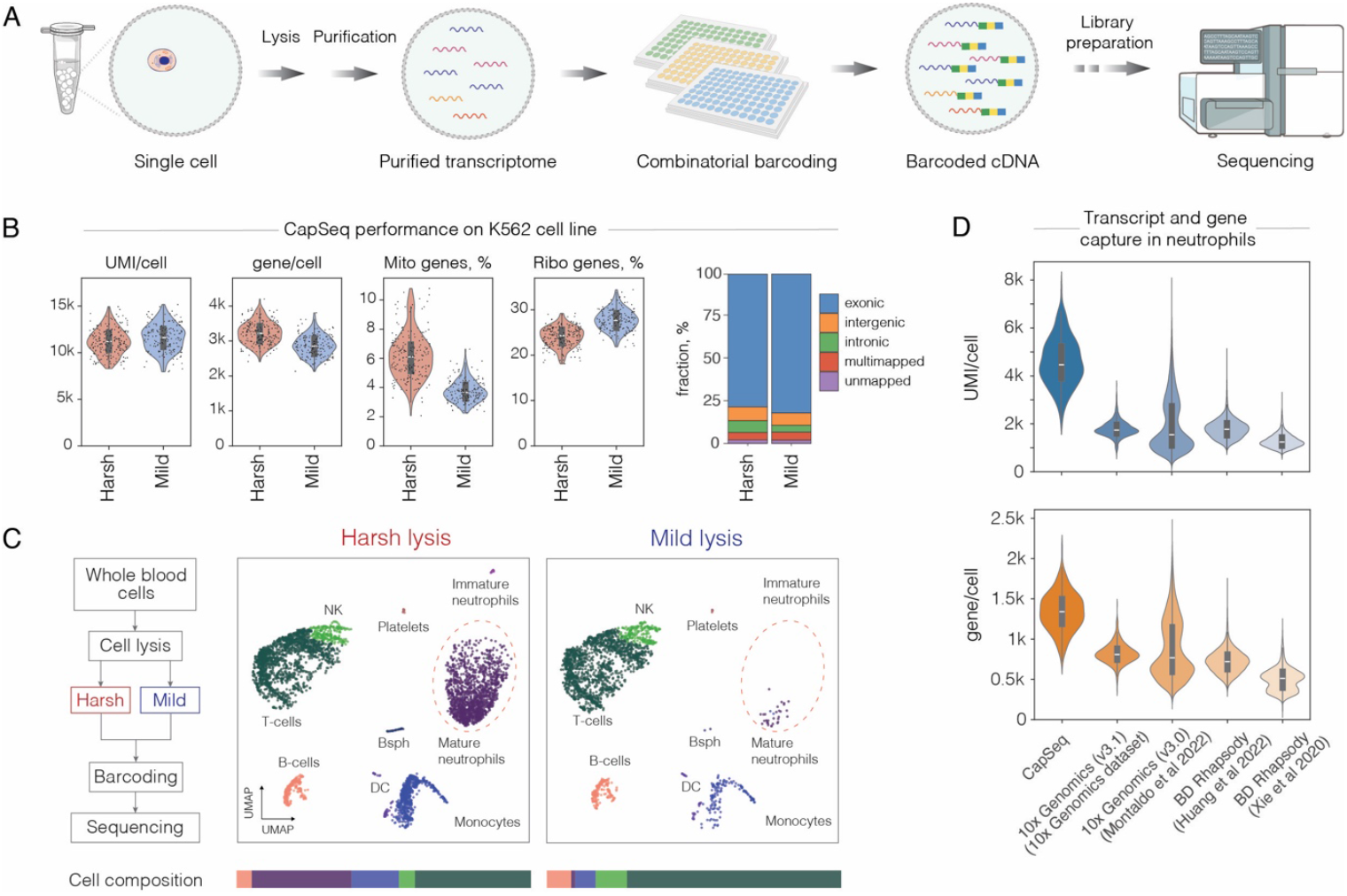
CapSeq performance. **A**) Schematics of capsule-based single-cell RNA-Seq (CapSeq) approach. **B**) CapSeq performance on K-562 cell line at a sequencing depth 20.000 reads per cell. **C**) Left: Flowchart detailing harsh and mild lysis workflow. Right: White blood cells from a healthy individual, prepared for scRNA-Seq under harsh and mild lysis conditions, projected on a UMAP. Note, the reduced recovery of neutrophils and basophils (Bsph) under mild lysis conditions. **D**) Comparative analysis of transcript and gene recovery in healthy human neutrophils with CapSeq, 10X Genomics (V3.1) and BD Rhapsody platforms (downsampled to 15’000 reads per cell).

We have optimized reverse transcription (RT) reaction, split-and-pool barcoding steps, and scRNA-seq library preparation aiming at improving transcript capture, barcoding efficiency, and reaction specificity (Methods). At first, we evaluated the performance of CapSeq by encapsulating K562 and NIH/3T3 cells and following split-and-pool barcoding we sequenced the libraries at a depth of 20’000 reads/cell. We obtained >10’000 transcripts and >3’000 genes per cell on average, with a mapping rate of 88.85% (**Figure 4B** and Figure S5). Compared to other scRNA-seq methods based on split-and-pool approach, CapSeq offers improved transcript capture (Figure S5B). To further assess CapSeq performance, we profiled white blood cells (WBCs) extracted from a healthy individual. We noticed that cell lysis conditions commonly used in droplet microfluidics and microtiter plate-based platforms failed to efficiently recover neutrophils; a serious limitation that has been reported previously (*28*). Compared to other immune cells, neutrophils are notoriously fragile, rich in nucleases, and carry the lowest amount of RNA (*29*) making them particularly challenging to capture and analyze. Only when we performed cell lysis with a strong denaturant (*i*.*e*., 4M GITC), we could successfully recover neutrophils and other hard-to-recover cell types such as basophils (**Figure 4C**). Therefore, CapSeq offers an immediate advantage for profiling the transcriptional state of highly sensitive and fragile cells that remain challenging to study using the existing scRNA-seq platforms.

After finding that harsh-lysis conditions are necessary to achieve efficient recovery of granulocytes, we conducted additional rounds of optimization aiming at improving UMI/gene capture in these and other immune cells. We noticed that a significant fraction of transcripts failed to incorporate PCR adapter by template switching (TS) reaction during cDNA synthesis despite our efforts to optimize RT reaction conditions (Figure S6). To rescue those cDNA molecules that failed to incorporate PCR adapter at the 5’ end we arrived at a protocol that includes a ligation step to attach a hairpin adapter (*30*) (Methods). This led to transcript recovery improvement over 3-fold (Figure S6), allowing us to achieve a significantly higher sensitivity for one of the most challenging cell types (*i*.*e*., neutrophils) than the best-performing approaches reported to-date (**Figure 4D**). These results illustrate that conducting scRNA-seq workflow, where each biochemical reaction is decoupled and conducted independently from a preceding one, can significantly improve transcript capture and recovery.

### Profiling hematopoietic disorders using CapSeq

To demonstrate the broader utility of CapSeq, we profiled WBCs extracted from patients suffering from hematopoietic disorders (Table S1-S3). Because mRNA in clinical samples is prone to fast degradation, which can lead to substantial biases in data analysis, we first safeguarded the transcriptional state of the cells by preserving them in methanol (Methods). In a series of independent experiments, we derived a protocol enabling reliable and reproducible recovery of WBCs following blood collection, preservation, rehydration, and single-cell mRNA sequencing (Methods). We confirmed that highly diverse cells of innate and adaptive immunity are recovered with proportions closely matching the results of a recent comprehensive study (*31*) (Figure S5D). Next, we performed scRNA-seq on 25 samples collected over the course of 6 months and obtained a comprehensive cell atlas consisting of 86,447 transcriptomes (**Figure 5**, Figure S7 and Methods). It included WBCs of acute myeloid leukemia (AML) patients (n = 8) and follow-up samples (n = 5) of the same individuals who developed neutropenia condition following chemotherapy treatment (Figure S7). Untreated healthy (n = 4) and granulocyte colony-stimulating factor (G-CSF) treated donors (n = 5) were also included in the study to serve as references for comparative analysis of granulocytes and other non-leukemic cells. The targeted DNA sequencing of AML samples revealed canonical mutations (*32*) in NPM1 (62% of patients), TET2 (62%), FLT3 (62%) and other leukemia-associated genes (Figure S7B and Table S4).

**Figure 5.**
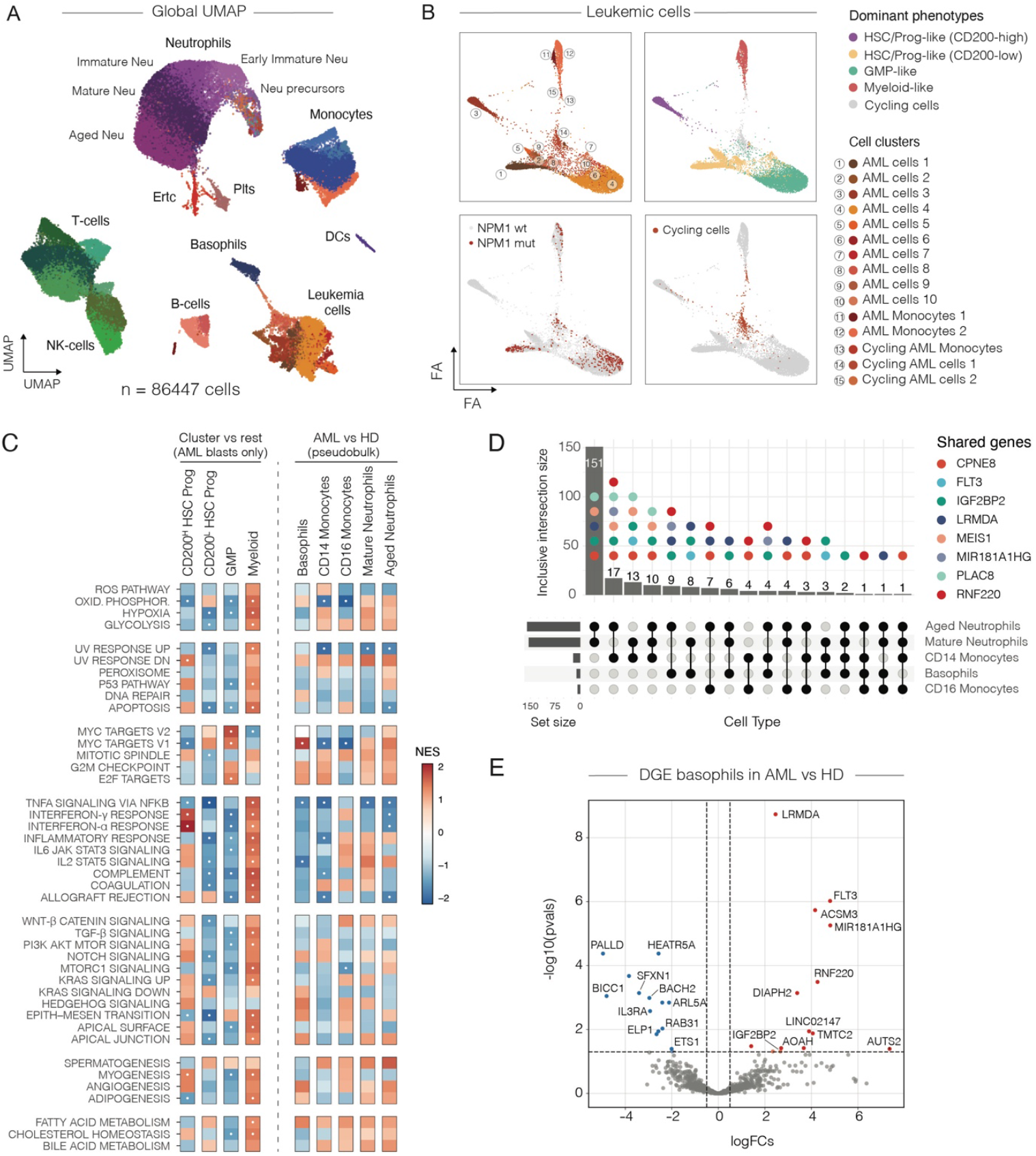
Transcriptional profiling of hematopoietic disorders using CapSeq. **A**) Single cell transcriptomes from 25 samples embedded on UMAP and colored according to annotated cell phenotypes. **B**) Force directed graph layout of the leukemic compartment. **C**) Gene set enrichment analysis of Hallmark pathways (MSigDB) was performed using ranked gene lists based on differential expression and statistical significance, comparing each AML cell cluster against all others (cluster vs. rest) or myeloid cell types between healthy donors and AML patients in pseudobulk analysis. Normalized enrichment score (NES) is represented by a color gradient, with blue indicating depletion (negative values) and red indicating enrichment (positive values). Statistically significant pathways (FDR < 0.05) are marked by a white dot. **D**) Upset plot showing shared differentially expressed genes among innate immune cell types when comparing AML patients and healthy donors. The bottom panel represents the intersecting cell types, while the top panel displays the size of the intersection (number of shared differentially expressed genes). Colored dots in the top panel indicate the most frequently shared genes across all intersections. **E**) A volcano plot illustrating all significantly differentially expressed genes in basophils from AML patients compared to healthy donors, with upregulated events shown in red and downregulated events in blue.

We applied size factor normalization (CP10K), log-normalization and principal component analysis (PCA) followed by batch integration (*33*) to project cells in the Uniform Manifold Approximation and Projection (UMAP) 2D space (**Figure 5A** and Figure S8). The unsupervised graph-based clustering (Leiden) followed by supervised marker gene analysis (*34, 35*) enabled identification of all major circulating cell types including a rich spectrum of neutrophil states reflecting the continuum of their maturation as well as sequential formation of primary (azurophilic), secondary (specific) and tertiary (gelatinase) granules (Figure S9). We also successfully recovered neutrophils from the AML patients despite their fraction being substantially depleted (Figures S9C and S9D). Furthermore, the biological processes important for neutrophil function were faithfully captured by CapSeq revealing that in the AML context these cells are struggling to mount an efficient immune response due to impaired chemo-sensing, migratory and degranulation activities (Figure S9G).

Consistent with minimal technical batch effects across samples, we observed low inter-patient heterogeneity (high Shannon entropy) among immune cells and high diversity in leukemic cells (Figure S10A). The leukemic compartment clustered into multiple and largely patient-specific cell populations (**Figures 5B**, Figures S7C, S7D, S8D, and Tables S5, S6), with cycling AML cells exhibiting the highest entropy among the patients (Figure S10B). Using established gene signatures (*36-38*) we assigned AML cells to three major hematopoietic phenotypes corresponding to HSC/Progenitor-like (which separated into CD200-high and CD200-low populations), GMP-like and myelomonocytic cells (**Figure 5B**, Figure S10C and S10D). The majority of immature cells showed exclusive upregulation of well-defined marker genes (Figure S10E and Tables S7, S8) associated with hematopoietic stem cell phenotype (e.g., *MEIS1 (38), SOX4, and ZNF521 (39, 40))*. Interestingly, expression of *HTR1F* was upregulated in HSC/Prog-like blasts but not in GMP-like progenitor cells, or myelomonocytic cells (Figure S8E and Table S9), indicating its potential use as a distinct marker for the HSC-like phenotype. In addition, blast and progenitor cells showed elevated expression of *CDK6*, a key regulator of the G1-S phase transition, and apoptosis suppressor *RNF220* (Figure S8E, Table S9). On the contrary, the myelomonocytic blasts, the presence of which was also confirmed by flow cytometry (Tables S10 and S11), were negative for cell proliferation markers (Figure S8E) and displayed elevated hypoxia signature (Figure S10F) suggesting that these cells might be circulating in a quiescent state (*41*). Gene set enrichment analysis (GSEA) further revealed that, when compared to other AML-related cells, myelomonocytes display a relatively strong pro-inflammatory phenotype, while HSC/Prog-like and GMP-like cells exhibited reduced inflammation (**Figure 5C**), along with upregulation of MYC and E2F targets, consistent with a proliferative response. Stratification of patients based on the dominant phenotypes correlated with ELN2022 risk classification (*42*); patients carrying GMP- or HSC/Prog-like leukemic cells showed favorable and adverse risk, respectively (Figure S10H), recapitulating similar findings in the bone marrow studies (*43, 44*).

### Mature innate immune cells in AML display a dysfunctional phenotype

Having confirmed that CapSeq faithfully captures relevant gene expression signatures in AML patients, next we sought to investigate the transcriptional landscape of mature innate immune cells. While basophils, neutrophils and monocytes execute different biological functions, we found that some genes in AML samples compared to healthy donors’ appear consistently upregulated in all or a fraction of these mature populations (**Figure 5D** and Tables S12-S16). The most shared genes across the granulocytes and monocytes (Table S17) were also expressed by blasts in our dataset (Table S18). Some of these genes (Table S17) remain poorly defined, yet others have important implications in cellular signaling, survival and differentiation, or are related to the general functioning of the innate immunity and development of AML disease (*44-49*).

The upregulation of *IGF2BP2, FLT3, MEIS1, PLAC8* and *MSRB3* in mature cells is particularly noteworthy given that these genes are typically associated with the progenitor/stem cell state of leukemic cells and are generally considered not to be expressed by differentiated cells. For instance, *IGF2BP2* was recently shown to be overexpressed in leukemia stem cells (LSC), promoting their self-renewal and AML development (*45*). The expression of FMS-like tyrosine kinase 3 (*FLT3*) is assumed to be limited to hematopoietic stem or progenitor cells, yet we observe expression of *FLT3* in mature basophils (**Figure 5E**) and neutrophils (Table S12), as well as differentiated CD14 monocytes (Table S17). Another example, *MEIS1*, a transcription factor thought to mark HSC/progenitor cells (*44*), when co-expressed with *HOXA9* (co-upregulated in CD14 monocytes from AML samples (Table S15)) denotes a specific monocytic-LSC program involved in progression and development of AML (*46*). On the same note, expression of *PLAC8*, also known as onzin, inhibits differentiation of AML cells (*47*), while *MSRB3* has been shown to induce a stem-like state (*48*). Finally, *FCGR1A* (Table S17), also known as CD64, albeit generally expressed by healthy monocytes, was shown to mark LSCs (*46*). Together, upregulation of this set of genes is an interesting finding that has not been reported to date, and potentially indicates altered, stem-like state of mature innate immune cells.

Dysregulation of intracellular signaling and membrane trafficking (*e*.*g*., increase in *CPNE8, LRMDA, EAF2, GAS7* expression) further supports notation that mature cells in AML exhibit a dysfunctional phenotype. Pathway enrichment analysis revealed significant TNFα signaling via NFκB downregulation across all the innate populations (except for CD16 monocytes, **Figure 5C** and Figure S9I). Interestingly, both CD16 and CD14 monocytes showed significant downregulation of OXPHOS, together with MYC targets V1 pathway, which was, conversely, upregulated in granulocytes. Furthermore, CD14 monocytes had impaired inflammatory response, while neutrophils were marked by diminished response to α and γ interferon signaling. GSEA analysis further highlighted corrupted basophil activation in AML patients, as prominent downregulation of IL2-STAT5, a pathway essential to basophil function, was observed (Figure S9J).

Taken together, our results indicate that in AML patients innate immune cells, but not lymphoid cells, share a strong signature of dysfunction and stemness.

### Single-cell RNA cytometry, sorting for regulatory RNAs expression and sequencing

A large fraction of differentially expressed genes in leukemic compartment encoded a variety of regulatory RNAs (**Figure 6A**), including highly abundant *MIR181A1HG, LNCAROD* and *LINC02147* (**Figure 6B**). In particular, our attention was drawn by upregulation of *MIR181A1HG* gene (**Figure 6C**), which encodes poorly characterized lncRNA that has been associated with multiple types of leukemias (*50, 51*) and is suspected to be involved in epigenetic regulation, as it produces at least two functional RNA molecules (*52, 53*). One of them is intronic small RNA miR-181a, which plays a key role in hematopoiesis (*54*) and AML pathogenesis (*55*). The other is a long non-coding RNA that is localized in the nucleus (*53*) and putatively interacts with several RNA-binding proteins, including the aforementioned *IGF2BP2* (Table S19), a gene commonly upregulated in the differentiated myeloid cells of AML patients (**Figure 5D**). In our dataset, *MIR181A1HG* was co-expressed with genes involved in hematopoiesis, cell development and differentiation (Figure S11).

**Figure 6.**
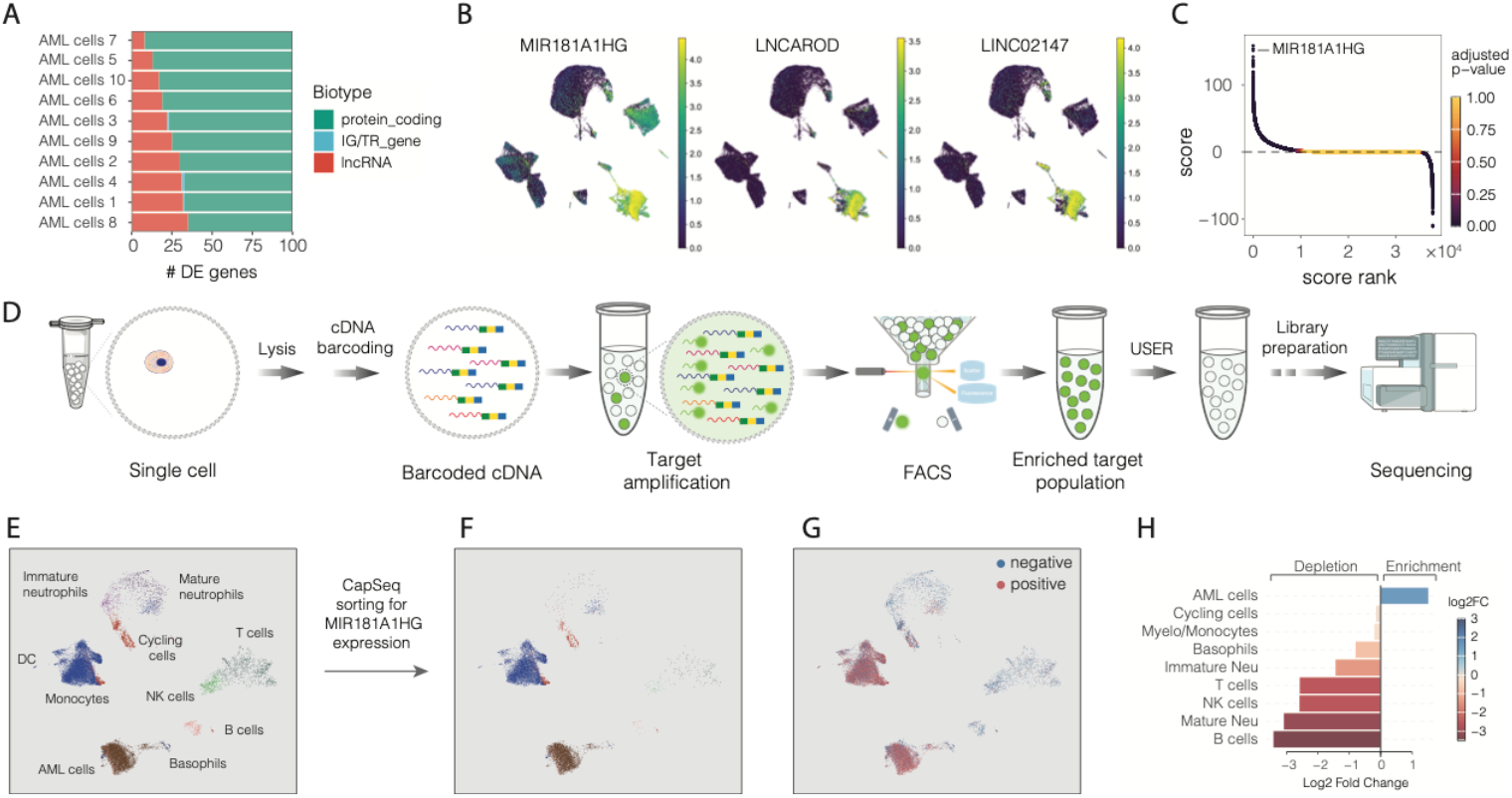
Single cell transcriptome enrichment by FACS for expression of a nucleic marker of interest. **A**) Gene biotype distribution among the top 100 differentially expressed genes in AML cells compared to all other cells. **B**) Single cell transcriptomes embedded on UMAP and colored for MIR181A1HG, LNCAROD and LINC02147 expression. **C**) Differentially expressed genes in AML cells, compared to all other cells, ranked by a combined score of differential expression and statistical significance. Each dot represents a gene and is colored by the adjusted P-value. Positive score values indicate upregulated genes, while negative values correspond to downregulated genes in AML blast cells. **D**) Experimental design for sorting and sequencing single cell transcriptomes based on the nucleic acid marker of interest. **E**) Single cell transcriptomes of 4 AML patients (# 2, 4, 6 and 7) embedded on UMAP and colored according to the cell type. **F**) Sorted single cell transcriptomes for expression of MIR181A1HG embedded on UMAP **G)** Positive and negative fractions of FACS-sorted transcriptomes based on MIR181A1HG expression **H**) Differential abundance analysis of cell types between the positive and negative fractions of FACS-sorted cells based on MIR181A1HG gene expression. Log2 fold change is shown on the x-axis and indicated by a color gradient, with blue representing depletion and red representing enrichment of a given cell type in the positive fraction of FACS-sorted cells.

To enrich the cells positive for *MIR181A1HG* we sought to combine CapSeq and single-cell RNA cytometry (*1*) and derive a general strategy allowing to acquire whole transcriptome sequencing data on single cells, FACS-sorted for a desirable nucleic acid marker (**Figure 6D**). At first, we confirmed that specificity of the target amplification from the barcoded cDNA library in SPCs remains high irrespectively of nucleic acid marker of interest (Figure S12A). In addition, to enable selective amplicon degradation and reduce potential carry-over contamination we conducted targeted-PCR in the presence of uracil (Methods), so that later we could selectively hydrolyze PCR amplicon with the USER™ enzyme mix (*56*). We validated two approaches for constructing scRNA-seq libraries, both of which produced the expected size and distribution of nucleic acid fragments (Figure S12B). Next, we encapsulated methanol-preserved WBCs from four AML patients, which included varying levels of *MIR181A1HG* expression across different cell types with a fraction of peripheral blasts ranging from 9.4% to 39% (Table S3). After the RT reaction followed by split-and-pool barcoding, we combined SPCs into one tube and subjected them to a PCR using fluorescently labeled primers targeting cDNA encoding *MIR181A1HG* (Methods). The resulting capsules clearly separated into fluorescent (having an amplicon) and non-fluorescent (lacking an amplicon) populations as they passed FACS instrument and were sorted to a positive fraction (**Figure 6E-G** and Figure S12C). Out of ∼17,000 sorted SPCs that were recovered, 15,000 were used for sequencing, out of which we obtained transcriptomes for 14,440 cells. Sequencing confirmed robust enrichment of *MIR181A1HG* positive cells and depletion of *MIR181A1HG* negative cells in all patient samples (Figure S13A and S13B). For example, sorted leukemic blasts were enriched in all patient samples, while other cell types were depleted (**Figure 6H**, Figure S13C, Table S20). Collectively, these results clearly show the broad utility of SPC technology beyond regular scRNA-seq applications as it supports nucleic acid detection, FACS-based isolation and sequencing of single cells based on the expression of RNA markers. This approach has a potential to expand the scope of single-cell assays that rely on isolation of phenotypes based on nucleic acid markers that lack appropriate antibodies and probes.

## Discussion

One of the major goals of single cell multi-omics methods is to provide a thorough and unbiased characterization of cellular genotype and phenotype both in health and disease, at scales ranging from several thousand to millions of cells and at a minimal cost. A large variety of single-cell -omics techniques developed to-date attempt to fulfill this goal (*16, 57*), yet the methodological and technological challenges of efficiently conducting complex molecular biology and biochemical workflows at high-throughput persist, restricting further applications.

Here, we describe a highly versatile and broadly applicable technology for single-cell biology research that is based on semi-permeable capsules, or SPCs. The SPCs are uniform, concentric compartments, having a liquid core enveloped by a thin, and tunable, semi-permeable shell (**Figure S2C, S2D**). Single cells and biomolecules are loaded in SPCs using high-throughput microfluidics, yet once SPCs are produced, all subsequent operations become microfluidics-free and can be carried out using standard laboratory pipettes and tubes, making the approach easy to adopt and implement. Being resilient, the SPCs sustain standard laboratory operations, freezing, thawing, thermocycling, sorting by FACS and different types of treatments with organic and inorganic solvents. Like hydrogel beads, SPCs support facile solvent exchange and compatibility with high-resolution imaging, yet in contrast to hydrogels the retention of nucleic acids in SPCs does not rely on physical embedding in a mesh, or hybridization to covalently conjugated oligonucleotides. This feature is particularly advantageous, as capture of nucleic acid fragments in a free (liquid) state is orders of magnitude more efficient as compared to capture on a solid support (*58, 59*). The unique and useful feature of SPCs described in this work is their complete and selective retention of nucleic acid fragments longer than 300 bp. (**Figure 1E**), while simultaneously allowing enzymes as large as 160 kDa to enter the SPC within minutes (**Figure 1F**) and interact with target RNA or DNA molecules. This permeability cut-off is therefore well-suited for a large variety of multi-omics workflows, and if needed might be further tuned by adjusting the concentration and modification degree of the monomers in the shell. Considering that accurate representation of phenotype and biological functions depends on faithful and efficient biomolecule capture, the SPCs are exceptionally well suited for the analysis of individual cells.

Another distinct advantage of SPCs is their exceptional biocompatibility. Single cells isolated within SPCs can be readily expanded into isogenic colonies over days – or even weeks (**Figure 3**) – thus overcoming a major limitation of droplet microfluidics technology. Cultivating encapsulated cells simply requires SPC dispersion in a cell culture flask. This straightforward experimental setup offers flexibility in designing complex cell-based assays that involve, for example, co-cultures of different cell types or rely on specific cell-to-cell communications via soluble factors. Importantly, releasing the cells is also elementary; exposing SPCs to collagenase enzyme breaks down the shell in a matter of minutes, enabling retrieval of single cells or microcolonies without damaging them, a prerequisite for functional cell-based screening assays.

Taking advantage of distinctive SPCs properties, we implemented a universal and highly customizable scRNA-seq approach, entitled CapSeq, based on combinatorial split-and-pool barcoding and sequencing. The high cell capture rate and superior sensitivity of CapSeq allowed us to profile the full diversity of peripheral blood cells of AML patients, including the extremely fragile innate immune cells that are particularly difficult to work with due to their minimal RNA content (*29*). CapSeq-enabled comprehensive depiction of cellular heterogeneity in AML revealed a shared dysfunctional, stem-like state in the mature innate immune cell compartment, including granulocytes. These results raise two equally likely hypotheses about the observed phenotypes: it may either reflect the shared cells’ response to the AML-induced microenvironment or suggest the enduring potential of AML blasts to differentiate into these altered mature phenotypes. Further mechanistic studies are needed to disentangle these scenarios.

Finally, we also report a transcriptome sorting method based on the expression of RNA markers, which we exemplify by FACS-based enrichment of AML cells upregulating lncRNA *MIR181A1HG*. To this day, FACS-based enrichment of nucleic acid markers remains challenging and mainly relies on fluorescence *in situ* hybridization (FISH) of chemically cross-linked cells (*60, 61*), or custom-built microfluidic platforms (*62*). Th use of SPCs provides a straightforward option for a high-throughput sorting of single transcriptomes based on a nucleic acid target of interest – a simplified and user-friendly option to enrich virtually any desirable cell population, without specific antibodies or FISH probes. Thus, compared to current scRNA-seq and FACS-based technologies, the SPCs offer large flexibility, establishing them as a versatile and scalable platform for next-generation single-cell assays.

## Acknowledgements

This work was supported by National Institutes of Health under grant R33 CA278392-01 and conducted as part of the execution of Project “Mission-driven Implementation of Science and Innovation Programmes” (No. 02-002-P-0001), funded by the Economic Revitalization and Resilience Enhancement Plan “New Generation Lithuania.” We thank Vilnius University Hospital Santaros Klinikos Biobank for providing biological samples and health information as well as patients who agreed to participate in this study. We also thank Genomics Core Facility at EMBL (Heidelberg, Germany) for their assistance. Authors are grateful to Monika Kirsnyte, Suji Kim, Mindaugas Sinis and Rapolas Zilionis for their valuable assistance.

## Author contributions

DB: development of CapSeq, implementation of single-cell preservation and rehydration protocol; DB and SN: single-cell RNA-seq experiments; development of FACS-based enrichment for nucleic acid markers; DB, JZ, SJ, KG, KS and LM: data analysis and interpretation; DB, SN, SJ, KG, GL prepared the images and graphs for figures; GL: development and implementation of semi-permeable capsules; GL, SN, VM: Evaluation of physical and biochemical properties of SPCs; GL: development of cell cultivation protocols in SPCs; GL: development of SPC-based single DNA molecule PCR, single-cell RT-PCR and whole genome amplification; VM: implementation of genome sequencing using SPCs; KS and LG: bioethics approval, biospecimen acquisition, and clinical study design; VK: optimization of sorting parameters; LM: conceptualization, study design, supervision and funding acquisition. LM: wrote the manuscript and prepared the figures; JZ, DB and LM: revised the manuscript; All the authors have proof-read, commented, and approved the final manuscript.

## Code and Data Availability

Sequencing data associated with this work is available at GEO accession GSExxxxx. The following publicly available datasets were used in this study: for the NIH/3T3 cell line benchmark Gene Expression Omnibus (GEO) at GSE110823 and GSE98561. For neutrophils benchmark, ArrayExpress under accession number E-MTAB-11188, NCBI repository PRJNA772373 and Gene Expression Omnibus (GEO) at GSE137540. Custom code and intermediate data files to reproduce the analyses supporting this work are available at https://github.com/mazutislab/capseq

