## Supplementary material for "High-throughput single cell -omics using semi-permeable capsules": Materials and Methods

### Cell lines

K562 (ATCC, CCl-243) and HEK293 (ATCC, CRL-1573) cells were grown in Iscove's Modified Dulbecco's Medium (IMDM, Gibco, 12440053) supplemented with 10% fetal bovine serum (FBS; Gibco, 10270106) and 1X penicillin-streptomycin (PS; Gibco, 15140122) at 5% $CO_2$ and 37 °C. NIH/3T3 (ATCC, CRL-1658) cells were grown in Dulbecco's Modified Eagle Medium (DMEM, Gibco, 61965026) supplemented with 10% fetal bovine serum and 1X penicillin-streptomycin (PS; Gibco, 15140122) at 5% $CO_2$ and 37 °C. Cells were harvested at a concentration of approximately $10^6$ cells/ml, transferred into 15 ml conical tubes, centrifuged at 300g for 5 minutes, and rinsed twice with ice-cold 1X Dulbecco's Phosphate Buffered Saline (DPBS; Gibco 14190144). Cell count and viability were determined using the hemocytometer and 0.2% trypan blue staining.

### Phase diagram of the mixing between GelMA-dextran

10% (w/v) Dextran solution in 1x PBS was equilibrated at room temperature, and 10% (w/v) GelMA solution in 1X DPBS was kept at 37 °C to prevent gelation. Both polymer solutions were combined at a given concentration at room temperature, and whether phase separation occurred was determined based on the turbidity of the solution.

### Scanning electron microscopy

Semi-permeable capsules (SPCs) were frozen and lyophilized overnight at -80 °C using a freeze drier. A small number of capsules was spread on double-sided carbon tape, magnetronically sputtered with chromium metal to increase the conductivity of the material and introduced into the scanning electron microscope (SEM) chamber for imaging. The imaging of SPCs was conducted in a dual-beam system of scanning electron microscope Helios Nanolab 650 equipped with a two-beam system with a Schottky-type field emission electron source and a gallium ion source. Cross-sections were made using a 30 keV focused Ga ion beam.

### DNA retention in SPCs

To determine the size of DNA fragments that can leave or be retained inside the SPCs, we loaded SPCs with GeneRuler Low Range DNA Ladder (Thermo Fisher Scientific (TFS), SM1191) comprising DNA fragments from 25 to 700 bp in size. The SPCs (150 µl) were resuspended in 5 ml of 1x DPBS supplemented with 0.1% F-68 and incubated at room temperature (22 °C) for 4 hours by inverting the tube every 30 min. At selected time points (0.5, 1, 2 and 4 h) 1 ml of SPCs suspension was retrieved from the tube and washed 3-times in 500 µl of 1x DPBS supplemented with 0.1% F-68. After a final wash, the SPCs pellet was dissolved by adding 2 mg/ml proteinase K (TFS, AM2548) for 10 minutes at 37 °C. The material released from dissolved SPCs was collected and analyzed on 3% agarose gel in 1x TAE buffer. The retention of DNA fragments was quantified by measuring the DNA band intensity on the agarose gel from three independent measurements and analyzed using Fiji software package.

### Permeability to proteins

To determine the diffusion kinetics of biomolecules and proteins into the core of SPCs we dispersed capsules in 1x PBS supplemented with either i) FITC dye, fluorescein isothiocyanate (0.38 kDa), FITC-labelled dextran (20 kDa), FITC-labelled bovine serum albumin (66 kDa) or FITC-labelled immunoglobulin G (160 kDa). We then recorded fluorescence intensity over time using a confocal microscope (Leica) and determined the half-time that it takes for the SPCs to reach half of its equilibrium fluorescence intensity. Based on the diffusion model for a hollow sphere with a thin shell the half-time can be expressed as $t_{1/2} \approx \frac{R^2}{6D}$, where *t* is the half-time for diffusion (s), *R* is the radius of the hollow sphere (m) and *D* is the effective diffusion coefficient of the analyte through the shell ($m^2$/s).

### Single-DNA molecule digital PCR

Single-DNA molecule PCR in capsules follows the same principles as digital droplet PCR (*1, 2*) and is based on the Poisson distribution of DNA molecules in SPCs; $P(X = k) = \frac{e^{-\lambda}}{k!} \lambda^k$ , where $P$ is the probability to have $k$ copies of DNA molecules per capsule, and $\lambda$ is the mean number of DNA molecules per capsule. We encapsulated pET29-GFP plasmid DNA (λ = 0.3) so that one SPC would receive, on average, no more than 1 copy of DNA. The PCR master mix (Platinum SuperFi, Invitrogen, 12358250) included 0.5 μM primers [pET T7 forward/reverse, see Table S21] that generate a 1510 bp amplicon. Following 35-cycles PCR (98°C for 30s; 35-cycles [98°C for 5s, 61°C for 10s and 72°C for 20s] and final extension at 72 °C for 5 min) we stained SPCs with DNA intercalating dye SYBR Green I (Invitrogen, S7563) and analyzed SPCs under fluorescence microscope. The SPCs containing amplified single DNA molecules during PCR became highly fluorescent upon staining and their fraction matching the expected $\lambda$ value. The SPCs that lacked DNA remained blank (non-fluorescent).

### Target specific RT-PCR in capsules

Single-cell RT-PCR was performed as described previously (*3*). Specifically, a mixture of cells K562 and HEK293 were loaded into SPCs (refer to section "Cell encapsulation" below) so that on average one SPC contains no more than 1 cell. The SPCs hosting single cells were suspended in GeneJET RNA Purification Kit Lysis Buffer (TFS, K0732) containing 40 mM DTT, washed once with fresh GenJET Lysis Buffer and incubated at room temperature for 5 minutes. Next, the SPCs were washed 5-times in 1 mL Washing Buffer (WB, 10 mM Tris-HCl [pH 7.5], 0.1% (v/v) Triton X-100) followed by treatment with 0.05 U/μl DNAse I enzyme (TFS, K2981) and 0.2 U/μl RNase Inhibitor (TFS, EO0381) at 37 °C for 20 minutes. Following gDNA depletion, the SPCs were rinsed 3-times in WB and subjected to a reverse transcription (RT) reaction in 200 μL reaction mixture comprising: 100 μL close-packaged SPCs, 1x RT Buffer (TFS, EP0751), 1x Oligo(dT)18 primer (TFS, SO131), 0.5 mM dNTP Mix (TFS, R0192), 5 U/μl Maxima H Minus Reverse Transcriptase (TFS, EP0751), 0.2 U/μl RiboLock RNase Inhibitor and incubated at 50 °C for 60 minutes. After cDNA synthesis, the SPCs were washed 3-times in WB and then subjected to target-specific PCR. The PCR was performed in 100 μl reaction volume comprising: 47 μl of close-packed SPCs, 50 μL of 2x Phire Tissue Direst PCR Master Mix, 0.5 μL 555-YAP-forward primer, 0.5 μL YAP-reverse primer, 0.5 μl 488-PTPRC-forward primer, 0.5 μL PTPRC-forward primer, 0.5 μL 647-ACTB-forward primer and 0.5 μM μL ACTB-forward (refer to Table S21 for primer sequences). All primers were at 100 μM concentration. The PCR was performed for 30-cycles (98 °C for 5 min followed by 30-cycles (98°C for 5s; 64°C for 5s, 72°C for 20s) and final extension at 72°C for 1 min). After the PCR, the SPCs were treated with Exonuclease I (NEB, M0293L) by adding 100U directly to post-PCR mix and incubated at 37 °C for 15 minutes. The fluorescence of SPCs was evaluated under epifluorescence microscope (Nikon Ti-Eclipse) using 10x (NA 0.45) objective. All centrifugation steps above were conducted at 2000g for 1-2 min.

### Single genome amplification in capsules

Single cells (K562) were loaded into SPCs (refer to section "Cell encapsulation" below) so that on average one SPC contains no more than 1 cell. The SPCs with K562 cells were suspended in GeneJET RNA Purification Kit Lysis Buffer (TFS, K0732) containing 40 mM DTT, washed once with fresh GenJET Lysis Buffer and incubated at room temperature for 5 minutes. Next, the SPCs were washed 5-times in 1 mL Washing Buffer (WB, 10 mM Tris-HCl [pH 7.5], 0.1% (v/v) Triton X-100) and subjected to multiple displacement amplification (MDA). The MDA was performed in 100 μL reaction volume by mixing 50 μL of close-packed SPCs with 50 μl of MDA reaction mix containing 1X Reaction Buffer (TFS, EP0091), 1 mM dNTP Mix (Invitrogen, 18427013), 25 μM Exo-Resistant Random Primer (TFS, SO181), 1 mM DTT (TFS, R0861) and 0.5 U/μL phi29 DNA Polymerase (TFS, EP0091). Whole genome amplification by MDA reaction was performed at 30 °C for 6 hours. After the MDA, capsules were rinsed once in 1 mL WB supplemented with 5 mM EDTA. Post-MDA capsules were stained with 5 μM SYTO 9 for 30 minutes at room temperature, then rinsed twice with 500 μl WB and analyzed under epifluorescence microscope (Nicon Ti-Eclipse) using 10x (NA 0.45) objective. All centrifugation steps above were conducted at 2000g for 1-2 min.

### Whole genome sequencing

As a proof-of-concept for whole genome sequencing, we isolated K562 cells in SPCs and lysed in 4M GITC (see harsh Cell lysis description below). After rinsing the capsules 10-times with 1 mL of ice-cold post-RT washing buffer (WB, 10 mM Tris-HCl [pH 7.0], 50 mM NaCl, 0.1% (w/v) Triton X-100, an Illumina sequencing library was prepared using the Collibri™ ES DNA Library Prep Kit for Illumina (Invitrogen, A38606024) following the manufacturer's instructions with some modifications. Specifically, the genome fragmentation reaction was assembled on ice by adding 5 µL of 10X Fragmentation and dA-tailing Buffer to 25 µL of packed capsules pre-mixed with 20 µL of 10 mM Tris-HCl (pH 7.5) buffer in a thin-wall PCR tube. The capsule suspension was mixed by pipetting up and down, and then 10 µL of 5X Fragmentation and dA-tailing Enzyme Mix was added to the reaction mixture. The fragmentation was performed for 20 minutes at 37 °C, followed by end-repair and dA-tailing for 10 minutes at 65 °C (with the lid set at 80 °C). Subsequently, the capsules were washed in 0.2 mL of ice-cold post-RT washing buffer, and a dual-indexed adaptor was ligated by adding 5 µL of 10X T4 DNA ligase buffer, 7.15 µL of dual-indexed adaptor and 1.5 µL of 5 U/µL T4 DNA ligase to 36.35 µL of capsule suspension in 10 mM Tris-HCl (pH 7.5) buffer. Ligation was performed for 1 hour at 22 °C, followed by inactivation of the enzyme for 10 minutes at 65 °C. Following ligation, the SPCs were dissolved by adding 0.01 mg/ml Proteinase K and the adaptor-ligated genomic DNA was size-selected for 350 bp inserts using DNA Cleanup Beads, and eluted in 21 µL of nuclease-free water. The purified DNA library was amplified by PCR using the following cycling conditions: an initial denaturation at 98 °C for 30 s, 12 cycles of 98 °C for 15 s, 60 °C for 30 s, and 72 °C for 30 s, followed by a final extension at 72 °C for 60 s. The PCR mix was prepared by combining 20 µL of the DNA library, 25 µL of 2X Library Amplification Master Mix, and 5 µL of Primer Mix. The amplified sequencing library was purified using DNA Cleanup Beads, (Invitrogen, A38606024), and eluted in 20 µL of nuclease-free water.

For the construction of an Illumina sequencing library from bulk genomic DNA, 150 ng of purified K562 genomic DNA was used following the guidelines of the Collibri™ ES DNA Library Prep Kit for Illumina (Invitrogen, A38606024). Specifically, genome fragmentation was conducted for 20 minutes at 37 °C, followed by end-repair and dA-tailing for 10 minutes at 65 °C. Next, a dual-indexed adaptor was ligated following size selection for 350 bp inserts. Subsequently, the adaptor-ligated product was amplified by PCR and purified using DNA Cleanup Beads.

The concentrations of gDNA libraries were measured using a Qubit 4 instrument, and the quality and size distribution of the libraries were assessed using the High Sensitivity DNA assay on an Agilent Bioanalyzer 2100. Sequencing was performed using the Illumina MiSeq Reagent Kit v2 Nano system with the following parameters: R1 (read 1) – 150 cycles; i7 – 8 cycles; and R2 (read 2) – 150 cycles.

### Single cell clonal expansion and spheroid formation in SPCs

Single cells were isolated in SPCs (refer to section "Cell encapsulation" below) using a microfluidics chip having rectangular channels of 40 µm height and a nozzle 40 µm wide (Figure S2A). The flow rates for IMDM cell culture medium carrying cells and supplemented with 15% dextran (500 kDa) was set at 100 µL/h, shell solution comprising 3% (w/v) gelatin methacrylate was set at 250 µL/h, and the carrier oil at 700 µL/h. Cell encapsulations were performed at 25-26°C for about 20-30 min. The SPCs were resuspended in 1x IMDM (Gibco, 12440053), containing 10% FBS (Gibco, 11573397) and 1x Penicillin-Streptomycin (Gibco, 15070063) and transferred to a cell culture flask, placed at 37 °C under

5% $CO_2$ atmosphere. The cell expansion was followed over several weeks, and at selected time points the SPCs were analyzed microscopically.

### Cultivation of microorganisms in capsules

*Escherichia coli* MG1655 and *Saccharomyces cerevisiae* were separately encapsulated at a limiting dilution. The *E. coli* strain was encapsulated in 40 µm size SPC, using a microfluidics chip 20 µm height and having a nozzle 20 µm wide. The yeasts were encapsulated in 55 µm size SPCs, using a microfluidics chip 40 µm height and having a nozzle 40 µm wide. The flow-rates for encapsulation of *E. coli* were: shell phase – 50 µL/h, core phase with bacteria– 20 µL/h and the carrier oil – 300 µL/h. Typical flow-rates used for encapsulating yeasts were: shell phase – 50 µL/h, core phase with cells – 40 µL/h and the carrier oil – 300 µL/h. The SPCs with microorganisms were rinsed twice in 1X PBS, containing 0.1 % Pluronic F-68, and then resuspended in 1 ml of culture media: LB-Miller containing 0.1 % (w/v) Pluronic F-68 for bacteria and YPD containing 0.1 % (w/v) Pluronic F-68 for yeasts. Finally, the SPCs (n ~ 100.000) were transferred to a 35 mm Petri dish (TFS, 130180), prefilled with 1 ml of corresponding culture media. Microcapsules with bacteria were incubated at 37 °C and recorded for 5 hours at 1-hour intervals. Microcapsules with yeasts were incubated at 30 °C and recorded for 15 hours at 2- to 4.5-hour intervals.

### Sample acquisition of human white blood cells

Human blood samples were collected from healthy and sick patients from the Vilnius University Hospital Santaros Klinikos Biobank (Vilnius, Lithuania) with a bioethics committee approval Nr. 2024/4-1583-1039. Samples were collected from a total of 8 AML patients, 8 neutropenia patients, 5 G-CSF stimulated donors, and 4 healthy donors.

### Red blood cells lysis

Red blood cells (RBC) lysis was performed in the following manner. Peripheral blood of fresh samples was collected into a K2EDTA vacutainer and stored on ice. Subsequently, 10 µL of 0.5 M EDTA (Invitrogen, cat. no. 15575-038) was added per 1 mL of peripheral blood. Next, 0.2 mL of peripheral blood was transferred to 2 mL Protein LoBind tube (Eppendorf, cat. no. 0030108132) and gently mixed with 1.8 mL of RBC lysis buffer (R7757, Sigma-Aldrich). Cells in RBC lysis buffer were incubated for 5 min at room temperature, gently inverting the tube every 5 s. Next, cells were centrifuged for 10 min at 300g and 4 °C using a swinging-bucket rotor. After discarding the supernatant, the pellet was gently resuspended with 2 mL RBC lysis buffer. The tube was incubated for 3 min at room temperature, gently inverting every 5 s. Then the cells were centrifuged for 5 min at 300g and 4 °C using a swinging-bucket rotor, and the supernatant was discarded. Additional wash in RBC lysis buffer was performed if the cell pellet displayed dark-red color. Finally, the pellet was resuspended to the final volume of 100 µL in ice-cold 1X DPBS.

### Cell fixation in methanol

Cells were dehydrated by slowly adding 900 µL of ice-cold methanol into the 100 µL of cell suspension in 2 mL Protein LoBind tube. To avoid cell clumping, the first 500 µL of the methanol was added drop by drop (3-5 drops per 1 s) and the remaining volume was added faster (in 1-2 s). After methanol was added, the tube was gently inverted 10-times, incubated on ice for 15-30 min and transferred at -20 °C (for a short period <3 months) or at -80 °C (for a longer period >3 months).

### Rehydration of methanol-fixed cells

The cells preserved in methanol were withdrawn from the freezer and incubated on ice for 15-30 min followed by centrifugation for 5 min at 1000g and 4 °C using a swinging-bucket rotor. Next, the methanol was discarded, and the cell pellet was resuspended in 1 mL of rehydration buffer (RB; 1X SSC, 2 M NaCl, 0.1% BSA). The resuspended cells were transferred to a 1.5 mL tube and centrifuged for 5 min at 1000g and 4 °C. After discarding the supernatant, the cells were subjected to an identical washing procedure using 1 mL of RB. A 10 µL aliquot of the cell suspension was used to determine the cell count on a hemocytometer. The cells were diluted with RB to achieve a final volume corresponding to 3 million cells per milliliter.

### Assessment of RNA quality after rehydration

RNA from white blood cells was extracted using in-house TRIzol reagent (*4*), following the TRIzol protocol (Thermo Fisher Scientific, cat. no. 15596026). The RNA integrity number (RIN) was assessed using the total RNA Pico Assay (Agilent Technologies, 5067-1513) with the Agilent Bioanalyzer 2100 instrument.

### CapSeq procedure

CapSeq procedure consists of the following steps: i) cell encapsulation, ii) cell lysis, iii) reverse transcription (BC1 addition), iv) 1$^{st}$ adapter ligation (BC2 addition), v) 2$^{nd}$ adapter ligation (BC3 addition) and vi) cDNA amplification. All the barcoding steps and the cDNA amplification step are performed inside the capsules. Primers and barcoding sequences could be found in **Tables S21-S25**. After cDNA amplification, the shell of the capsules is dissolved, barcoded cDNA is released, and prepared for sequencing on the Illumina platform. A step-by-step protocol is provided in "Supplementary Protocol 1" file.

### Cell encapsulation

Single cell encapsulation was performed as described previously, (*3*) with some minor modifications. The stock solution of Dextran (Sigma, 31392) was prepared in 1X DPBS at a 30% (w/w) concentration, the stock gelatin methacryloyl (GelMA; Sigma, 900496) solution was prepared in 1X DPBS at a 10% (w/w) concentration. The working solution of 15% (w/v) Dextran with cells was prepared by mixing the equal volumes of 30% (w/w) Dextran solution and cell suspension, achieving a final dilution of ~1.5 million cells per 1 ml. The working solution of 4.5% (w/v) GelMA was prepared by diluting pre-heated (37 °C for 30 min) stock GelMA solution. Cell encapsulation was performed on a microfluidics platform Onyx (Atrandi Biosciences, MHN-ONYX1) using a 40 µm deep microfluidic device (Figure S2A). The flow rates were set at 100 µL/h for cell suspension containing dextran, 200 µL/h for GelMA solution and 700 µL/h for carrier oil. The resulting emulsion was collected in a 1.5 mL tube. To prevent premature GelMA solidification, the cell encapsulation was conducted at 26 ˚C.

Upon completion of cell encapsulation, the emulsion was transferred to 4 °C for 30 min, to solidify the shell. Continuing procedures on ice, SPCs having physically cross-linked shell were recovered from the carrier oil by first adding 500 µL of ice-cold Capsule Washing Buffer (CWB; 1X DPBS, 0.1% (v/v) Pluronic F-68) and then releasing the capsules with Emulsion Breaker (Atrandi Biosciences, MON-EB1). After a 5-minute incubation on ice, the capsules were transferred to a new 1.5 mL Protein LoBind tube, mixed with 500 µL of CWB, washed twice and then supplemented with 0.1 % (w/v) LAP (Sigma-Aldrich, 900889). The tube with capsules was exposed to low-energy light for 20 seconds using the Light Exposure Device (Atrandi Biosciences, MHT-LAS1) to obtain covalently crosslinked shells. The resulting SPCs were rinsed 2-times in CWB and used for subsequent biochemical reactions. All the washing and centrifugation steps were conducted at 300g for 1 minute at 4 °C.

### Cell lysis

Cell lysis was performed under harsh or mild lysis conditions.
For the harsh lysis, 200-300 µL of packed SPCs were mixed with 1 mL of harsh lysis buffer (4 M Guanidinium thiocyanate, 55 mM Tris-HCl [pH 7.0], 25 mM EDTA, 3 % (w/v) Triton X-100 and 40 mM DTT) by pipetting, and immediately centrifuged. Subsequently, the supernatant was aspirated and replaced with 1 mL of fresh harsh lysis buffer. The mixture was then incubated at room temperature (21°C) for 5 minutes. Following incubation, the SPCs were washed in 1 mL of ice-cold pre-RT washing

buffer (0.1X SSC, 50 mM NaCl, 0.1% (w/v) Triton X-100). When lysis was performed in multiple tubes, SPCs were pooled into a single 2 mL Protein LoBind tube after the last washing step.
When conducting mild lysis, the SPCs were resuspended to a final volume of 500 µL with CWB. Subsequently, 500 µL of mild lysis buffer (0.2X SSC, 0.6 % IGEPAL CA-630, 0.2 U/µL RiboLock RNase Inhibitor, 80 mM DTT) was added and mixed by pipetting. The mixture was then incubated at room temperature (21°C) for 5 minutes. Following incubation, the capsules were rinsed 5-times in 1 mL of cold pre-RT washing buffer. The centrifugation steps were carried out at 300g for 1 minute at 4 ˚C and the supernatant was aspirated.

### Barcoding plates preparation

The plates with barcoding oligonucleotides were prepared in the final working concentration of 50 µM for reverse transcription (RT) reaction (BC1 stock plate), 12 µM for 1$^{st}$ ligation (BC2 stock plate) and 10 µM for 2$^{nd}$ ligation (BC3 stock plate). To prepare the BC1, BC2 and BC3 working plates, 2 µL of each oligo from the stock plates were transferred using a multichannel pipette into new 96-well plates, with each oligo being allocated to rows A-H. Following this, the working plates were sealed and spun down at 1000g for 1 minute at 4 °C using a swinging bucket rotor. Finally, the prepared working plates were stored at -20 °C for future use.

### Reverse transcription – BC1 addition

The first barcoding step involves RT reaction. The SPCs were resuspended in cold pre-RT washing buffer in a 2 mL Protein LoBind tube to achieve a final volume of 1056 µL. Next, RT reagents were added into the tube in the following order: 422.4 µL of 5X RT Buffer (Thermo Fisher Scientific (TFS), EP0753), 100.32 µL nuclease-free water, 105.6 µL of dNTPs (10 mM dNTP mix; TFS, R0191), 5.28 µL of RNase Inhibitor (40 U/µL RiboLock RNase Inhibitor; TFS, EO0382), 105.6 µL of Template Switching Oligo (0.5 mM TSO – /5Biosg/AAGCAGTGGTATCAACGCAGAGTACATrGrGrG; HPLC purification, IDT) and 105.6 µL of Maxima Reverse Transcriptase (200 U/µL Maxima H Minus Reverse Transcriptase; TFS, EP0753). The RT master solution was mixed by pipetting, and 18 µl distributed into each well of a 96-well BC1 working plate containing 2 µl of 50 µM BC1 oligonucleotides (Table S22). The RT reaction was performed by incubating the BC1 working plate at 42 °C for 150 minutes without heat inactivation. Once the RT reaction was completed, the SPCs were pooled and mixed with 25 mM of EDTA. Subsequently, the pooled SPCs were washed 12-times with 1 mL of ice-cold post-RT washing buffer (10 mM Tris-HCl [pH 7.0], 50 mM NaCl, 0.1% (w/v) Triton X-100). The centrifugation steps were carried out at 300g for 1 minute at 4 ˚C.

### 1$^{st}$ ligation – BC2 addition

The second barcode (BC2) was added via ligation. Following RT barcoding step, the SPCs were resuspended in ice-cold post-RT washing buffer in a 2 mL Protein LoBind tube to achieve a final volume of 1056 µL. Next, ligation reagents were added into the tube in the following order: 211.2 µL of 10X T4 DNA Ligase Buffer (TFS, EL0013) 598.4 µL nuclease-free water and 35.2 µL of T4 DNA Ligase (30 U/µL T4 DNA Ligase; TFS, EL0013). The ligation reaction mixture was well mixed with SPCs and distributed by 18 µL into each well of a 96-well BC2 working plate containing 2 µl of 12 µM BC2 oligonucleotides (Table S23) using a multichannel pipette. After mixing the content of wells, the plate was incubated at 22 °C for 60 minutes. Next, the SPCs were pooled, and ligation reaction terminated with addition of 25 mM EDTA. Subsequently, the SPC suspension was rinsed 12-times with 1 mL of ice-cold post-RT washing buffer (10 mM Tris-HCl (pH 7.0), 50 mM NaCl, 0.1% (w/v) Triton X-100). The centrifugation steps involved 300g for 1 minute at 4 ˚C.

### 2$^{nd}$ ligation – BC3 addition

The third barcode (BC3) was added via ligation in the same manner as BC2 addition using BC3 working plate (Table S24) instead of BC2 working plate.

### Hairpin ligation

After completing split-and-pool barcoding of cDNA, the RNA in RNA:cDNA hybrid was digested using RNases mix, purified and ligated to a hairpin adapter. To digest RNA:cDNA hybrid, SPCs were typically resuspended in ice-cold post-RT washing buffer to achieve a final volume of 1000 µL. Next, RNA digestion reagents were added into the tube in the following order: 200 µL of 10X RNase H Reaction Buffer (NEB, M0297S), 720 µL nuclease-free water, 40 µL of RNase H (5 U/µL; NEB, M0297S), 40 µL of RNase A/T1 Mix (2 mg/mL; TFS, EN0551). Reaction mixture was well mixed by pipetting and RNA digestion reaction performed in a dry bath at 37 °C for 30 minutes. After RNA digestion was completed, SPCs were washed with 1 mL of ice-cold post-RT washing buffer and resuspended in ice-cold post-RT washing buffer up to a volume of 1000 µL Next, ligation reagents were added into the tube in the following order: 200 µL of 10X T4 DNA Ligase Buffer (TFS, EL0013) 636.7 µL nuclease-free water, 30 µL of ATP (100 mM; TFS, R0441), 100 µL of Hairpin-TSO oligo (0.1 mM Hairpin-TSO oligo [Table S21]), 33.3 µL of T4 DNA Ligase (30 U/µL T4 DNA Ligase; TFS, EL0013). The hairpin ligation reaction solution was well mixed by pipetting, split by 100 µL into multiple PCR tubes, and incubated at 16 °C for 16 hours. Following the ligation reaction, the SPCs were pooled into a single 2 mL Protein LoBind tube, washed several times with 1 mL of ice-cold post-RT washing buffer.

### Preparing SPC samples with an equal number of cells

To ensure reproducibility and consistency of the CapSeq workflow, we aimed to perform biochemical reactions on an equal number of encapsulated cells. For that purpose, 10 µL of diluted SPCs suspension (~20-100 SPCs/µl) were stained with SYBR™ Green I Nucleic Acid Gel Stain (Invitrogen, S7563), dispersed on a hemocytometer and evaluated under fluorescence microscope. After determining the number of fluorescent SPCs, we obtain cell occupancy values based on which we divide SPCs into equal parts so that each part (tube) would contain approximately 10'000 cells.

### cDNA library amplification

A suspension of SPCs containing 10'000 cells (as above) was centrifuged using a swinging-bucket centrifuge rotor at 2000g for 1 minute at 4 °C. Supernatant was aspirated and SPCs were resuspended in post-RT washing buffer to achieve a final volume of 48.8 µL. Next, 0.6 µL of forward cDNA amplification primer (100 µM CapSeq-REV) and 0.6 µL of reverse cDNA amplification primer (100 µM CapSeq-FWD) were added and mixed with 50 µL of 2X KAPA HiFi HotStart ReadyMix (Kapa Biosystems, KK2602). The cDNA was amplified within the SPCs by 15-cycles of PCR: 95 °C for 3 min, 15-cycles [98 °C for 20 s, 65 °C for 15 s, 72 °C for 5 min], and 72 °C for 10 min. After PCR was completed, SPCs were transferred from PCR tubes into separate 1.5 mL DNA LoBind tubes (Eppendorf, 0030108051) and washed 5-times with 1 mL of ice-cold post-RT washing buffer. The centrifugation steps were carried out at 300g for 1 minute at 4 ˚C. The excess supernatant was aspirated, and SPCs were resuspended in ice-cold post-RT washing buffer to achieve a final volume of 98 µL.

### Release of the capsule material

To dissolve SPCs and release the amplified cDNA material, 2 µL of 20 mg/mL Proteinase K (TFS, EO0491) was added into the 98 µL of capsule suspension and incubated at 37 °C for 10 min.

### Purification of amplified cDNA library

Each aliquot of the cDNA library released from the dissolved SPCs was diluted to a final volume of 400 µL by adding 300 µL of nuclease-free water. Subsequently, the cDNA library was purified with GeneJET PCR Purification Kit (TFS, K0702) and eluted in 100 µL with Elution Buffer. Following column purification step, the amplified cDNA was further purified using 0.6X SPRIselect beads (Beckman Coulter, B23318) and eluted in 20 µL with nuclease-free water. Next, the DNA concentration was measured using Qubit

4 instrument (TFS, Q33238) and Quant-iT™ 1X dsDNA HS Assay Kit (TFS, Q33232). Finally, the quality and size distribution of the purified cDNA was assessed with High Sensitivity DNA assay (Agilent Technologies, 5067-4626) on Agilent Bioanalyzer 2100.

### Illumina sequencing library preparation – indexing PCR

Illumina sequencing libraries were prepared for each amplified cDNA aliquot with NEBNext® Ultra™ II FS DNA Library Prep Kit for Illumina (NEB. E7805S) following the manual, with minor modifications. Specifically, 5-10 ng of amplified cDNA was used for fragmentation in the volume of 13 µL. Fragmentation reaction was assembled on ice by adding 3.5 µL of NEBNext® Ultra™ II FS Reaction Buffer followed by 1 µL of NEBNext® Ultra™ II FS Enzyme Mix. Reaction mix (17.5 µL) was vortexed and spun-down for 3-5 seconds. The fragmentation was performed for 8 minutes at 37 °C, followed by end-repair and dA tailing for 30 min at 65 °C (lid at 75 °C). Fragmented DNA library was purified with 0.6X-0.8X double-size selection using SPRIselect beads and eluted in 17.5 µL with nuclease-free water. Then, 1.5 µM ligation adapter was prepared in advance by mixing 1.5 µL of forward ligation primer (Table S21) and 1.5 µL of reverse ligation primer (Table S21) with 97 µL of water. To create dsDNA, the solution was heated to 95 °C for 2 min (lid at 100 °C) and cooled at room temperature for 5 min and used for subsequent ligation as a ligation adapter. Purified fragmented cDNA was mixed with 15 µL of NEBNext® Ultra™ II Ligation Master Mix, 1.25 µL of ligation adapter and 0.5 µL of NEBNext® Ligation Enhancer. Reaction solution was well mixed by pipetting and incubated for 15 min at 20 °C (lid set at 37 °C). The ligation product was diluted to 100 µL with nuclease-free water, followed by a 0.8X SPRI size selection procedure and eluted in 40 µL with nuclease-free water. Finally, the ligation product was amplified by 12-cycles of PCR: 95 ºC for 3 min, 12-cycles [98 ºC for 20 s, 54 °C for 30 s, 72 ºC for 20 s], and 72 ºC for 60 s. The PCR mixture included 20 µL of ligation product, 2.5 µL of forward indexing primer (Table S25), 2.5 µL of reverse indexing primer (Table S25) and 25 µL of 2X KAPA HiFi HotStart ReadyMix. Amplified libraries were purified with 0.6X-0.8X double-size selection using SPRIselect beads and eluted in 20 µL with nuclease-free water. Next, DNA concentration was measured using Qubit 4 instrument, and the quality and size distribution of libraries were assessed with High Sensitivity DNA assay on Agilent Bioanalyzer 2100. Sequencing was performed on Illumina NextSeq 2000 system, and the sequencing parameters were: R1 (read1) – 43 cycles; i7 – 6 cycles, R2 (read2) – 89 cycles (depending on instrument and sequencing reagent kit), at a depth of >10’000 reads per cell.

### Raw sequencing data pre-processing

The STARsolo wrapper pipeline (https://github.com/jsimonas/solo-in-drops) was used to process the sequencing data and to obtain cell × gene count matrices. Shortly, barcode ligation linker sequences were removed from the read1 and read1 was reassembled to have a structure of BC3[8nt]BC2[10nt]BC1[8nt] UMI[8nt]. Next, STAR (*5*) (version 2.7.10a) was run with the following parameters: --soloUMIdedup Exact, --soloFeatures GeneFull, --soloType CB_UMI_Complex, --soloCBposition 0_0_0_7 0_8_0_17 0_18_0_25, --soloUMIposition 0_26_0_33, --soloCBmatchWLtype EditDist_2. The reads were aligned to either the human GRCh38 genome (GENCODE v41) or the mouse GRCm39 genome (GENCODE M30).

### Data downsampling

For methods comparison scRNA-seq data was downsampled according to the scripts developed by Juzenas et al., (*6*). Each cell barcode in the bam file was downsampled to 15000 reads/barcode using "*downsample_by_coverage.sh"* script.

### CapSeq optimization

To identify the optimal enzymatic conditions for the CapSeq procedure, multiple conditions were tested (Figure S6) and sequenced. The sequencing data was initially downsampled using the *"downsample_by_coverage.sh"* script (*6*). Each cell barcode in the BAM file was downsampled to 2000 or 4000 reads per barcode. Next, sequencing saturation was calculated by further downsampling the newly generated BAM files to a given proportion of raw reads (5%, 10%, 20%, 30%, 40%, 50%, 60%, 70%, 80%, 90%, and 100%) using the *"downsample_by_fraction.sh"* script (*6*), which correspond to the number of detected UMIs. To reduce noise from outliers, the top 5% and bottom 5% of cells were excluded from the downsampling data. The sequencing saturation data points were then fitted using a generalized saturation function ($Y = Y_{max} \cdot (X / (k + X))$), and the UMI/cell at 80% sequencing saturation was extrapolated for each cell. Cells were grouped by their experimental condition. To determine the median UMI/cell, bootstrapping (100 realizations) was applied for each condition. The median and 95% confidence intervals were calculated from bootstrapping generated distributions. Finally, the median UMI count for each condition was normalized to the median value of the reference condition.

### scRNA-seq data analysis, quality control and doublet removal

Single-cell RNA sequencing data analysis was performed in Python using scanpy (1.10.2) package (*7*). Data analysis notebooks are provided in https://github.com/mazutislab. The lowest quality cells were removed from the cell x gene count matrices. Specifically, UMIs/cell thresholds were set manually for each sample (encapsulation run) by visually inspecting the total counts distribution. For the species-mixing experiment, the cut-off was set at 1000 UMIs/cell. For white blood cell samples, the initial cutoff was set at 200 UMIs/cell and then adjusted to 300. In all samples, cells with mitochondrial gene count fraction higher than 20% were filtered out. Doublets were removed using Scrublet (0.2.3) package (*8*). Each encapsulation run was treated separately with *expected_doublet_*rate parameter set from 0.01 to 0.05 depending on the observed doublet rate. The *min_gene_variability_pctl* parameter was chosen (typically from 80-95) to give the best-defined bimodal distribution based on the histogram of doublet scores generated by Scrublet. The threshold for doublet detection was set manually by inspecting the minimum between the two modes of the simulated doublet histogram. The other Scrublet parameters were left as default. Cells with a *doublet_scores* greater than 0.1-0.2 (depending on encapsulation run) were called as doublets and excluded for further analysis. Also, a second round of doublets removal was performed after initial clustering and annotation. Briefly, cells were re-clustered at high resolution using Leiden algorithm, and doublet scores and proportions of predicted doublets in each cluster were analyzed on a scatter plot. Clusters exhibiting high doublet scores, high proportions and having no unique gene expression in a cluster, were identified, and discarded.

### UMAP, clustering and annotation

UMAP construction was carried out as follows: gene expression was normalized to 10000 counts per cell. Mitochondrial and ribosomal genes were excluded, and an additional gene filter was applied to remove low-quality or noisy genes. Specifically, genes expressed in fewer than five cells with expression counts (counts per ten thousand) below 10 were removed. Subsequently, a log-transformation was performed. Highly variable genes were identified using the *scanpy.pp.highly_variable_genes* function with the *"seurat_v3"* flavor and the top 2500 genes selected. These genes were z-score scaled, and principal component analysis (PCA) was applied. Batch correction was performed using the Harmony tool (*9*). An adjacency graph was constructed with the *scanpy.pp.neighbors* function (n_neighbors = 30), and UMAP embedding was generated using the *scanpy.tl.umap* function (min_dist = 0.25). Initial clustering was performed using *scanpy.sc.tl.leiden* graph-based Leiden clustering (resolution = 1.4). Marker genes from the CellTypist database (*10*) for immune cell types were utilized to annotate cells. Next, low-complexity clusters were identified and removed. Then, second round of clustering was conducted on the filtered dataset using the graph-based Leiden clustering (resolution = 2.5), with additional subclustering within individual clusters to achieve finer resolution. To refine subtype identification for Neutrophils, T cells, and AML cells, the subsets of cells were re-clustered, and re-annotation was performed. A total of 86447 cells were retained for analysis.

### Differential gene expression

Before conducting differential gene expression analysis, genes expressed in fewer than five cells with expression counts (counts per ten thousand) below 10 were removed. Differential gene expression analysis was carried out for each cell type using the Wilcoxon rank-sum test with Benjamini-Hochberg correction. Genes were considered differentially expressed if the adjusted p-value was $< 0.05$ and the log2 fold change (log2FC) was $> 1$. The results were then sorted by log2FC. Differential gene expression analysis was performed separately for cell types identified in the global UMAP, for a subset of AML blasts, and across AML phenotypes.

### Phenotype assignment to AML blasts

To assign phenotypes to AML blasts (HSC/Prog-like, GMP-like, and Myeloid-like), we utilized signature genes identified by Van Galen et al. (*11*). First, all AML blasts were subsetted, and lower quality cells (<2000 counts) were additionally excluded. A force-directed atlas was then constructed using *scanpy.tl.draw_graph*. For each phenotype, the top 30 signature genes were selected, and their average expression was calculated for each cell. These expression values were subsequently z-score scaled. Cells were assigned to a phenotype based on the highest average expression among the phenotypic signatures. Finally, the predominant phenotype for each cell type was assigned based on the phenotype that was shared across the majority of cells within a given cluster.

### Pseudobulk analysis

Pseudobulk analysis was performed using decoupler package (*12*). Low-quality samples were filtered based on the number of cells (min_cells = 10) and the total sum of counts (min_counts = 1000). Additionally, noisy expressed genes were filtered using *decoupler.plot_filter_by_expr* function (min_count = 10, min_total_count = 15). Once pseudobulk profiles were generated, differential expression analysis was performed by comparing the gene expression of cells from diseased patients against healthy controls using DESeq2 (*13*) framework. Also, decoupler package was utilized to perform pathway activity inference using PROGENy (*14*) and transcription factor activity inference using CollecTRI (*15*). Over-representation analysis (ORA) was performed on the list of differentially expressed genes ($p < 0.05$) compared against gene sets from MSigDB database (*16*).

### Gene expression program inference by non-negative matrix factorization

To extract gene programs from single-cell expression data, non-negative matrix factorization (NMF) was applied on CP10k-normalized (10'000 total counts per cell) and log-transformed expression matrix using scikit-learn (v1.4.2) implementation of the NMF algorithm in Python. More explicitly, the matrix was decomposed into 25 latent factors (components), resulting in: (i) a cell-by-component matrix (W), representing the activity of each component in individual cells; and (ii) a component-by-gene matrix (H), representing the contribution of each gene to the components. To identify the most contributing genes for each component, knee point detection was performed using the KneeLocator function from the kneed (v0.8.5) Python package, enabling the removal of low-scoring genes. The remaining genes were used for the functional annotation of each gene expression program (component) through overrepresentation analysis (ORA), implemented in the decoupler package (*12*), using Gene Ontology Biological Process terms (*17*).

### Functional analysis

For the functional analysis of single-cell data, the decoupler package (*12*) was utilized, which provides a comprehensive collection of computational methods. To infer pathway activity PROGENy was used.

Gene Set Enrichment Analysis (GSEA) and Gene Set Variation Analysis (GSVA) were performed using *run_gsea* and *run_gsva* functions, respectively.

### RNA velocity analysis

For the RNA velocity analysis, neutrophils from this study and neutrophils from Montaldo et al. (*18*) were selected. To obtain spliced/unspliced gene count matrices, datasets were processed with STAR (*5*) (version 2.7.10a) with --soloFeatures Gene Velocyto parameters. First, UMAP representation was constructed using scanpy package (*7*) (version 1.10.3). In short, cells with at least 1000 counts and genes with at least 10 counts and found at least in 5 cells were kept for the analysis. The expression matrix was normalized, log-transformed, and 2500 highly variable genes were selected using *scanpy.highly_variable_genes()* function with flavor argument set to "cell_ranger". Next, PCA was performed, data was integrated using Harmony algorithm (*9*), and a k-neighbor graph (k = 100) was constructed. The graph was used to build UMAP with min_dist = 0.3. To obtain RNA velocity graphs, scvelo package (*19*) (version 0.3.3) was used. Data preprocessing was performed as described above for spliced and unspliced matrices until k-neighbor graph (k = 100) was constructed. Next, moments were computed using *scvelo.pp.moments*, which were used to estimate RNA velocity with *scvelo.tl.velocity* with mode parameter set to "stochastic". The estimated velocities were used to construct a velocity graph with *scvelo.tl.velocity* function, and the graph was used to embed RNA velocities into the UMAP constructed beforehand.

### NMP1 mutation identification

The NPM1 mutation was identified by analyzing BAM files using samtools v1.17 (*20*). Reads mapping to the GRCh38 reference genome at chromosome 5 (position 171,410,527–171,410,564) were extracted using the following command:
samtools view -h $bam "GRCh38_chr5:171410527-171410564" | grep "TCTCTGTCTGGCAG" | awk '{for(i=12;i<=NF;i++) if($i ~ /^CB:Z/) print substr($i, 6)}'. The command filtered reads containing the characteristic NPM1 insertion sequence ("TCTCTGTCTGGCAG") and retrieves the corresponding cell barcodes from the BAM file.

### Estimation of Shannon Entropy for cell types

To quantify variability in patient representation within each cell type, Shannon entropy was calculated. For each cell type, 100 cells were randomly sampled 100 times. In each iteration, the number of cells belonging to each patient was counted to compute the probability distribution. Shannon entropy was then calculated using the formula: $H = -\sum_{i=1}^{n} p_i \cdot log\,(p_i)$, where $p_i$ represents the proportion of cells from patient *i* in the sample.

### Targeted RNA molecule amplification

Targeted RNA molecule amplification was performed following a method described previously (*3*) with some modifications. First, SPCs were generated on a microfluidic co-flow device with a nozzle height of 20 µm and nozzle diameter of 20 µm (Atrandi Biosciences, MCN-C2). The flow rates were set to 500 µL/h for the oil phase, 40 µL/h for the GelMA phase, and 20 µL/h for the cell suspension in dextran phase. The cell concentration was ~ 6M/mL. The SPCs containing cells were treated under the harsh lysis condition and single cell transcriptomes barcoded following Supplementary Protocol 1. Next, SPCs were pooled and strained twice through the 40 µm mesh size strainer (TFS, 22-363-547) and subjected to hairpin adapter ligation. Then, SPCs were treated with 0.2 M NaOH containing 0.1 M EDTA at 95 °C for 5 min, followed by five washes with post-RT washing buffer and two additional washes with the post-RT washing buffer containing no NaCl. Targeted cDNA amplification was performed in two subsequent PCR reactions. PCR reaction mix consisted of 0.02 U/µL Phusion U Hot Start DNA polymerase (TFS, F555), 0.2 mM each dNTP (TFS, R0192), 0.2 mM dUTP (TFS, R0133), forward and reverse *MIR181A1HG* primers (MIR181A1HG_fwd1_ATTO488 and MIR181A1HG_rev1, refer to Table S21) of 0.5 µM concentration, and 1X Phusion HF reaction buffer, with the SPC suspension comprising half of the reaction volume. The first PCR was carried out for 20 cycles: 98 °C for 3 min, 20 cycles [98 °C for 5 s, 64 °C for 30 s, 72 °C for 30 s], and 72 °C for 5 min. Then, capsules were washed 3-times with post-RT washing buffer and twice with the post-RT washing buffer containing no NaCl. The second PCR was carried out under identical conditions but for 30 cycles. After target amplification, capsules were rinsed once with harsh lysis buffer and 5-times with post-RT washing buffer.

### Fluorescence microscopy of targeted amplicon in SPCs

Fluorescence images of SPCs were recorded using a Nikon Eclipse Ti microscope equipped with a Nikon Digital Sight DS-U3 camera. Imaging was performed with a FITC filter, an exposure time of 500 ms, and an analog gain of 7.6. For excitation, a CoolLED pE-300 illumination system was used, with the blue light source intensity set to 60%.

### FACS-based sorting of single cells transcriptomes based on target RNA

The sorting of SPCs was performed using a BD FACS Aria III instrument with BD FACSDiva V8.0.1 software (BD Biosciences, Franklin Lakes, NJ, USA). Standard optical filters and mirror configurations were used for the detection of scatter and fluorescence signals. For ATTO488 probe detection 488 nm laser with 530/30 BP filter was used. To operate the sorter, BD FACSFlow sheath fluid was used, and manufacturer's standard startup, cleaning, and QC procedures were performed. Setup & Tracking (CS&T) beads and drop calibration using AccuDrop beads (BD Biosciences) were applied before sorting. Sorting was carried out using a 100-μm nozzle with sheath pressure at 20 psi and plate voltage 2500 V with a slight increase in neighboring drop charge. Frequency was set to 22.5 kHz, drop delay was set to 21.26 (ticks of the system clock). The ND 1.5 filter was utilized, and sorting was performed in Purity mode, where Yield and Purity masks were set to the 32, and Phase mask to the 0, with sorting event count below 1000 events per second. SPCs were sorted into 15 mL falcon-type tubes prefilled with 10 mL of the post-RT washing buffer. SPCs were sorted based on side scatter (SSC) and fluorescence signal in the 530/30 channel and two-way sorting of ATTO488 positive and negative capsules was performed with 95 and 98% efficiency, respectively.

### Capsules recovery after sorting and amplicon removal

To recover SPCs after sorting, samples were spun down in a pre-cooled 4°C centrifuge (Eppendorf, 5428000610) for 5 min at 2200g. The SPC pellet was then used for further enzymatic processing. The SPCs from the positive and negative both fractions were treated with the USER® (NEB, M5505S) to remove the PCR amplicon. Treatment with the USER® was carried out in a 50 μL reaction, containing up to 25 μL of packed SPCs, and incubated at 37 °C for 30 min. An aliquot before USER® treatment was taken for imaging and DNA analysis with the Agilent Bioanalyzer 2100 instrument. After USER® treatment, the cDNA libraries were amplified by 15-cycles of PCR, released from SPCs, and fragmented for Illumina sequencing as described above.

### Calculation of sorting enrichment

To determine the sorting enrichment or depletion of different cell populations, a differential abundance (DA) analysis with edgeR (*21*) was performed on paired positive and negative sorting fractions from the same patients. A quasi-likelihood model was fit using a ~0 + sample_id + sorting_fraction model formula. Statistical significance of differences in cell population abundance between positive and negative fractions was assessed using empirical Bayes quasi-likelihood F-tests. Also, sorting enrichment was determined by calculating the fold enrichment between positively and negatively sorted fractions using the formula: Fold Enrichment = (Fraction of cell type in the positively sorted fraction) / (Fraction cell type in the negatively sorted fraction).

### Overlaying FACS data on Global UMAP Coordinates

To overlay FACS-sorted transcriptomes onto the global UMAP coordinates, the post-FACS dataset was first processed independently using the same approach as described above with some modifications. An adjacency graph was constructed with the scanpy.pp.neighbors function (n_neighbors = 50), and UMAP embedding was generated using the scanpy.tl.umap function (min_dist = 0.15). Next, the AnnData objects of the global UMAP and FACS datasets were concatenated, normalized to 10,000 counts per cell, and log-transformed. The same highly variable genes were selected as for the global UMAP, and subsequently z-score scaled. PCA embeddings were computed manually by taking the dot product of the principal component (PC) loadings from the global UMAP and the gene expression matrix of the concatenated AnnData object. Batch correction was performed using Harmony (*9*). Finally, an adjacency graph was constructed, and UMAP embeddings were generated for the concatenated AnnData using the same parameters as the global UMAP (n_neighbors = 30, min_dist = 0.25, respectively).

### Benchmark analysis

For the NIH/3T3 cell line benchmark, the following datasets were used: Gene Expression Omnibus (GEO) at GSE110823 (*22*) and GSE98561 (*23*). For Neutrophils benchmark: ArrayExpress under accession number E-MTAB-11188 (*18*), NCBI repository PRJNA772373 (*24*), and Gene Expression Omnibus (GEO) at GSE137540 (*25*).
