## Supplementary Information for "High-throughput single cell -omics using semi-permeable capsules"

### Supplementary Figures

**Supplementary Figure S1. Phase separation diagram. 2**

**Supplementary Figure S2. Generation of semi-permeable capsules. 3**

**Supplementary Figure S3. Capsule stability in organic and inorganic solvents and enzymatic hydrolysis. 4**

**Supplementary Figure S4. Structural properties and biological applications of semi-permeable capsules. 5**

**Supplementary Figure S5. Validation of CapSeq technology and cell preservation in methanol. 6**

**Supplementary Figure S6. CapSeq optimization to improve UMI capture. 7**

**Supplementary Figure S7. Clinical timelines of AML patients, treatment regimens, and responses over time. 8**

**Supplementary Figure S8. Single-cell RNA sequencing of AML, Neutropenia, G-CSF-treated, and healthy donors. 9**

**Supplementary Figure S9. Single-cell transcriptomic profiling of neutrophil maturation across AML patients, G-CSF-treated donors, and healthy individuals. 10**

**Supplementary Figure S10. Phenotypic diversity and classification of AML blasts. 11**

**Supplementary Figure S11. Long non-coding RNA MIR181A1HG is one of the major contributors to AML cell type-specific gene expression programs associated with cell differentiation, development, and activation. 12**

**Supplementary Figure S12. Experimental workflows for single-cell RNA cytometry and sorting based on targeted RNA expression. 13**

**Supplementary Figure S13. Cell type distribution and MIR181A1HG expression before and after capsules sorting on global UMAP coordinates. 14**

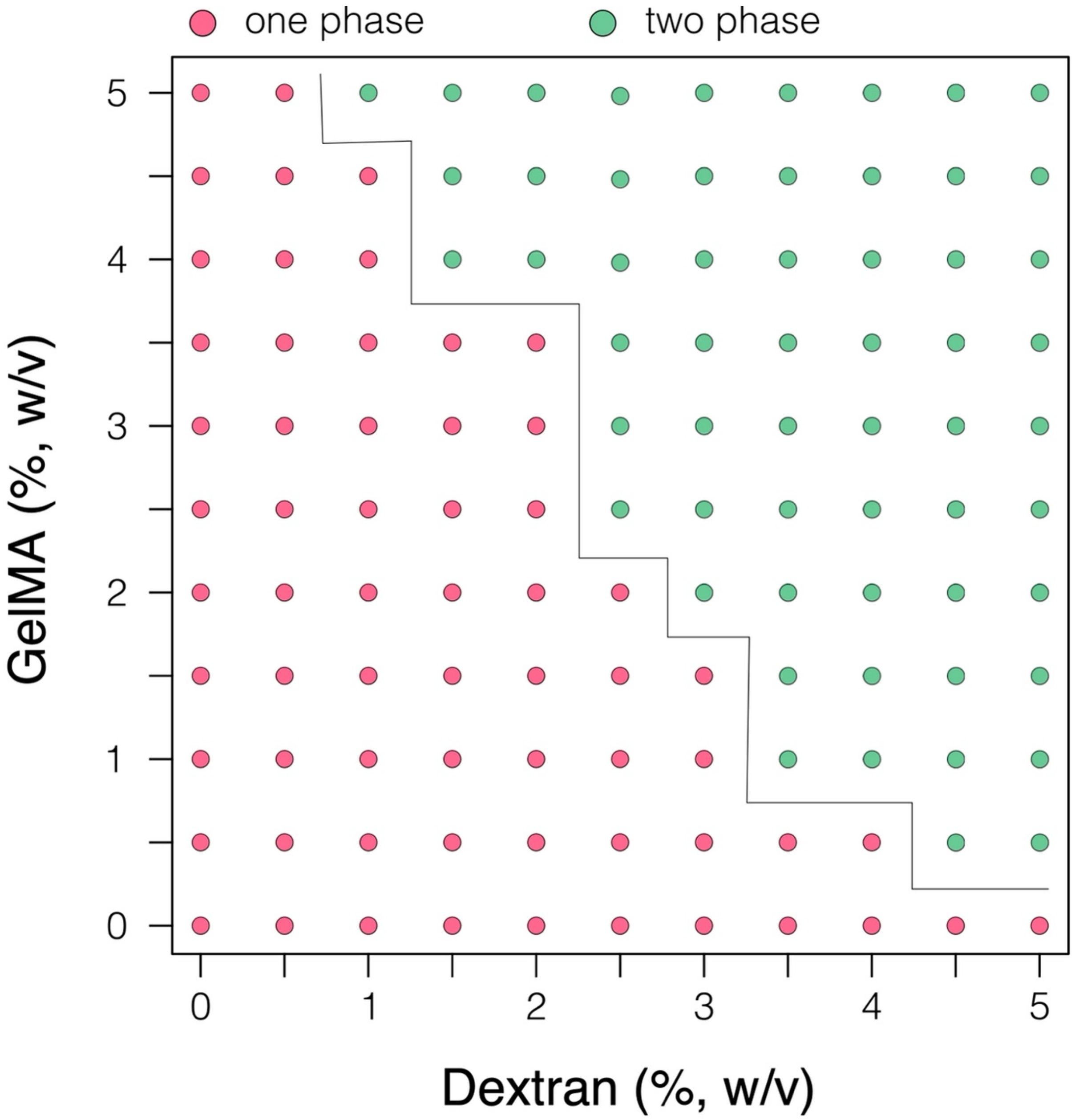


**Supplementary Figure S1. Phase separation diagram.** Two liquids, comprising dextran 500 kDa and gelatin-methacrylate (GelMA) were evaluated for liquid-liquid phase separation at 22 ºC in a phosphate buffered saline [pH 7.0]. The concentrations of GelMA and Dextran are indicated on Y- and X-axis, respectively.

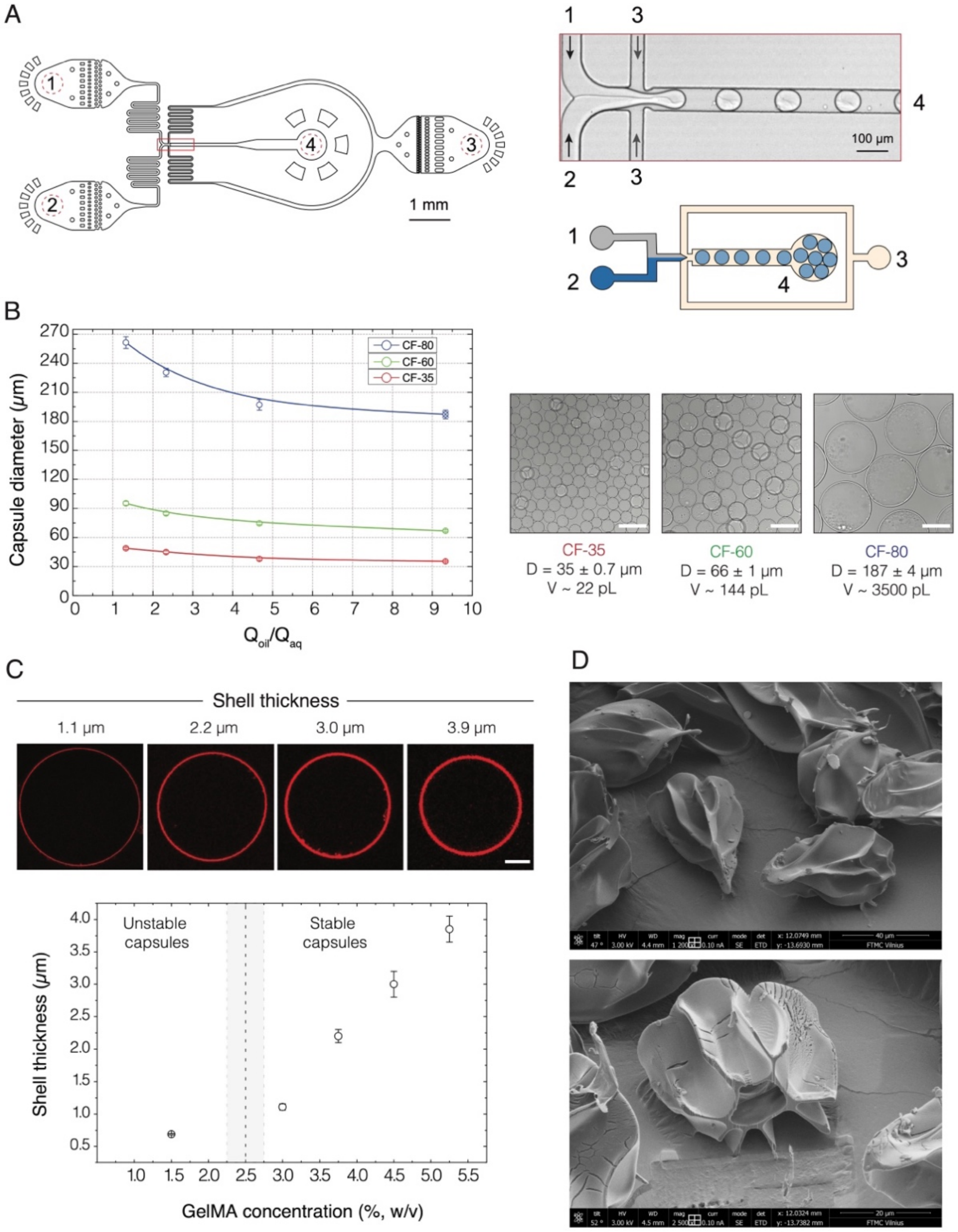


**Supplementary Figure S2. Generation of semi-permeable capsules. (A)** The design and operation of the microfluidics device. The microfluidics device contains two aqueous inlets (#1 and #2) for injecting core and shell forming solutions, carrier oil inlet (#3) and an outlet for collecting the emulsion (#4). The still image captures droplet generation. Scale bar, 100 µm. The droplets are generated at a flow-focusing junction, collected at the outlet, and are converted to SPCs using a two-step process as detailed in Materials and Methods section. **(B)** Capsules' size as a function of the flow rate ratio of carrier oil vs aqueous phase. Three microfluidic devices having 35 µm (red), 60 µm (green) and 80 µm (blue) deep microchannels were used to generate SPCs of different diameter while keeping a fixed flow ratio of carrier oil ($Q_{oil}$) vs aqueous phase ($Q_{aq}$). For example, for $Q_{oil}/Q_{aq}$ = 1.33 we used flow rates 50 µl/h for dextran solution, 100 µl/h for GelMA solution and 200 µl/h for carrier oil. For $Q_{oil}/Q_{aq}$ = 9.33 we used flow rates 25 µl/h for dextran solution, 50 µl/h for GelMA solution and 700 µl/h for carrier oil. The digital micrographs show SPCs obtained with different microfluidics chip at a flow ratio $Q_{oil}/Q_{aq}$ = 9.33. Scale bars, 100 µm. D – SPC outer diameter, V – SPC volume, CF-xx indicates microfluidics device where 'xx' indicate the depth of microfluidics chip in µm. **(C)** The generation of SPCs having a shell of tunable thickness. Top row shows confocal microscopy images of single SPCs with shells from 1.1 to 3.9 µm thick. Scale bar 20 µm. The graph below shows the shell thickness as a function of GelMA concentration (i.e., initial GelMA concentration that was loaded into a microfluidics chip). **(D)** Scanning electron microscopy of SPCs.

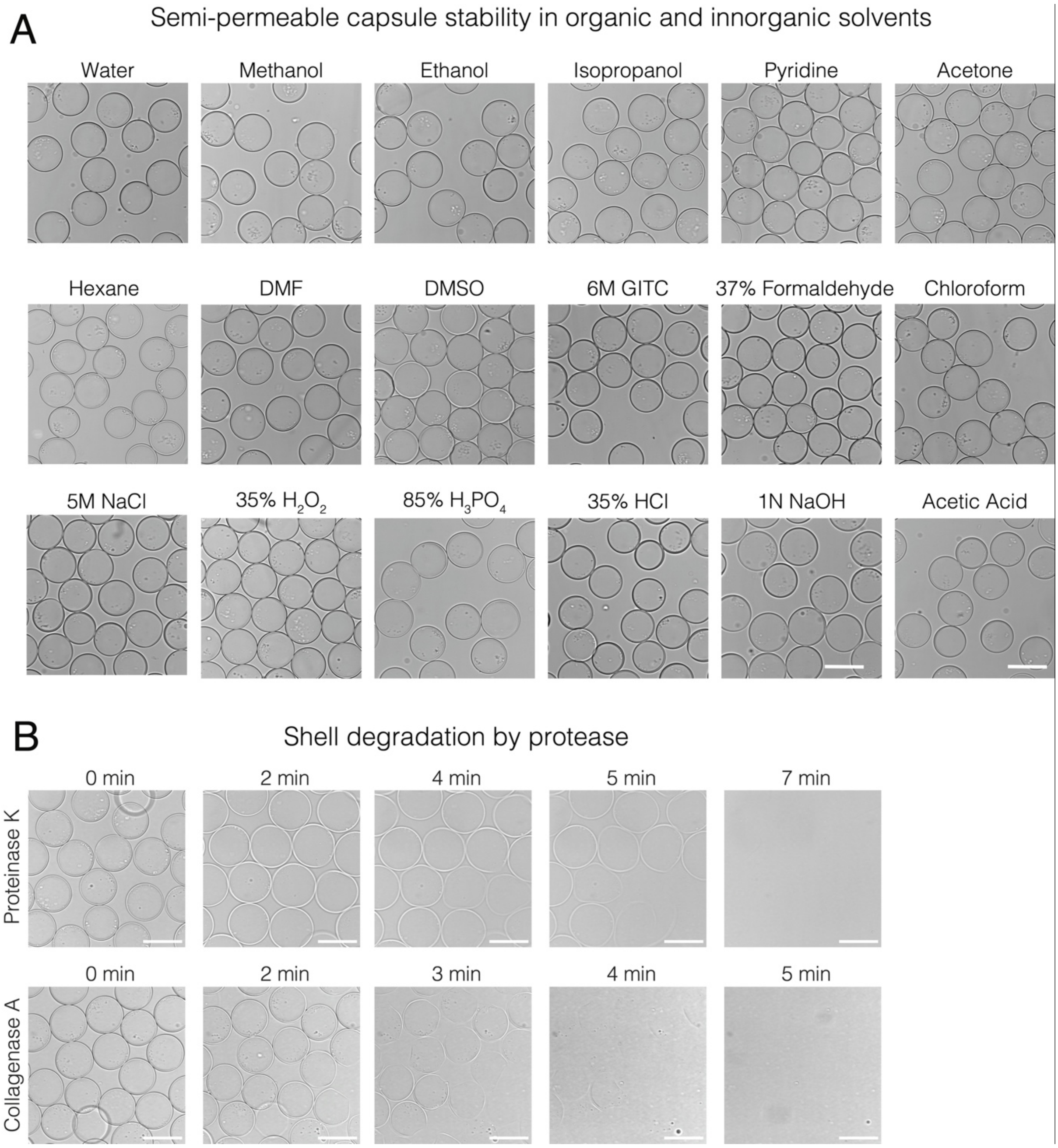


**Supplementary Figure S3. Capsule stability in organic and inorganic solvents and enzymatic hydrolysis.** **(A)** SPC stability in different organic and inorganic solvents. Capsules were exposed to different solvents, including water, methanol, ethanol, isopropanol, pyridine, acetone, hexane, dimethylformamide (DMF), dimethyl sulfoxide (DMSO), 6M guanidinium thiocyanate (GITC), 37% formaldehyde, chloroform, 5M NaCl, 35% $H_2O_2$, 85% $H_3PO_4$, 35% HCl, 1N NaOH, and acetic acid. Capsules were incubated for 30 min at 22 ºC. **(B)** Enzymatic hydrolysis of the SPCs shell. The final amounts of enzymes in the solution were 0.2 mg/ml and 1.0 mg/ml for Proteinase K and Collagenase A, respectively. Reaction was performed at 22 ºC. Scale bars, 100 µm.

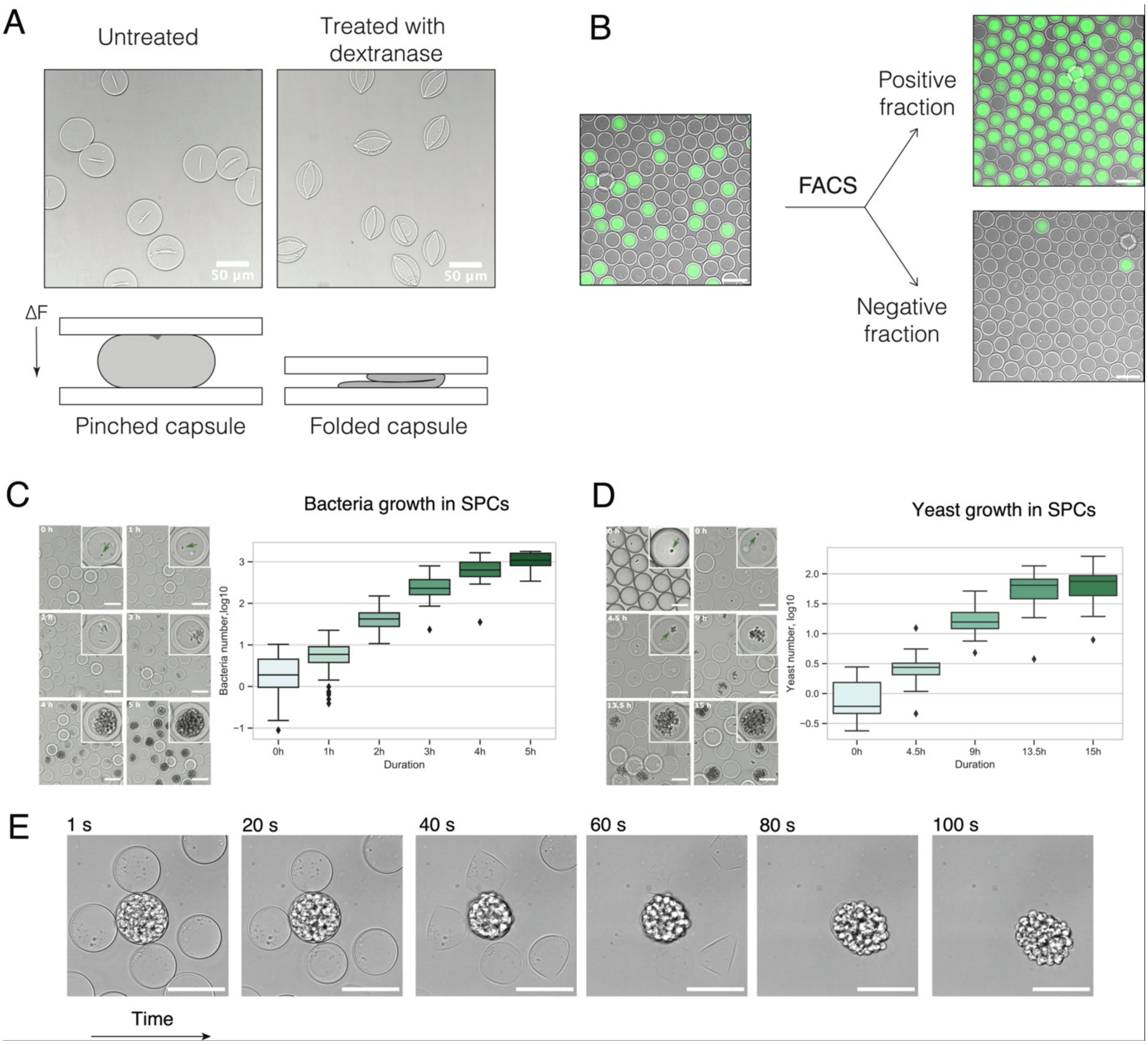


**Supplementary Figure S4. Structural properties and biological applications of semi-permeable capsules. (A)** Mechanical deformation of SPCs. Dextran within the SPC core prevents shell collapse under compression. Untreated capsules retain their spherical shape under compression (pinched capsule), while treatment with dextranase causes structural collapse, leading to folded morphology. Scale bars 50 µm. **(B)** FACS-based SPCs sorting. SPCs are compatible with high-throughput sorting using conventional FACS instruments. **(C)** Growth of bacteria and **(D)** yeast within SPCs. Time-lapse microscopy images show microorganisms proliferation inside SPCs into isogenic microcolonies. Box plots show the microorganisms number over time. **(E)** A time-lapse of an enzymatic hydrolysis of the SPCs shell using collagenase, allowing gentle and fast release of encapsulated cells. **(F)** Gel electrophoresis analysis of nucleic acids extracted from SPCs. Lane M: DNA marker. Lanes 1–4: DNA marker extracted from SPCs after multiple washing steps. DNA fragments larger than 300 bp are fully recovered from SPCs.

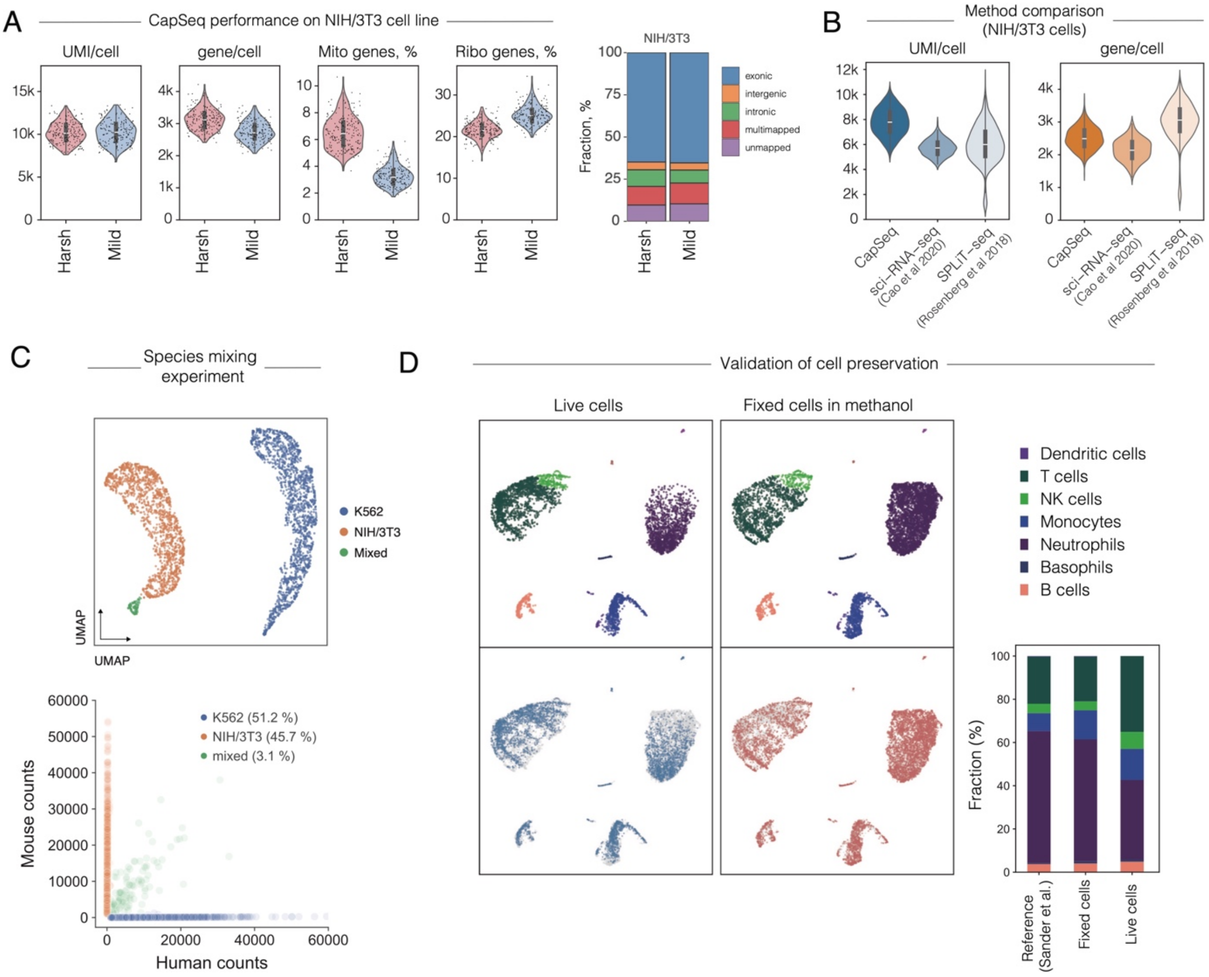


**Supplementary Figure S5. Validation of CapSeq technology and cell preservation in methanol. (A)** CapSeq performance on NIH/3T3 cells under Harsh and Mild lysis conditions. Violin plots show the number of UMIs per cell, genes per cell, mitochondrial gene percentage, and ribosomal gene percentage. The bar plot indicates the distribution of exonic, intronic, intergenic, multimapped, and unmapped reads. **(B)** Comparison of CapSeq with other plate-based scRNA-seq methods. Violin plots show the number of detected UMIs per cell and genes per cell. **(C)** Species mixing experiment of human (K562) and mouse (NIH/3T3) cells. UMAP visualization shows distinct clustering of K562 and NIH/3T3 cells, with a small proportion of mixed cells. The scatter plot bellow shows the human (x-axis) versus mouse (y-axis) counts per cell. **(D)** Validation of white blood cells preservation by methanol fixation. UMAP plots compare cell type clustering between live and methanol-fixed cells. The bar graph on the right compares the relative proportions of different immune cell types across reference data, fixed, and live cells.

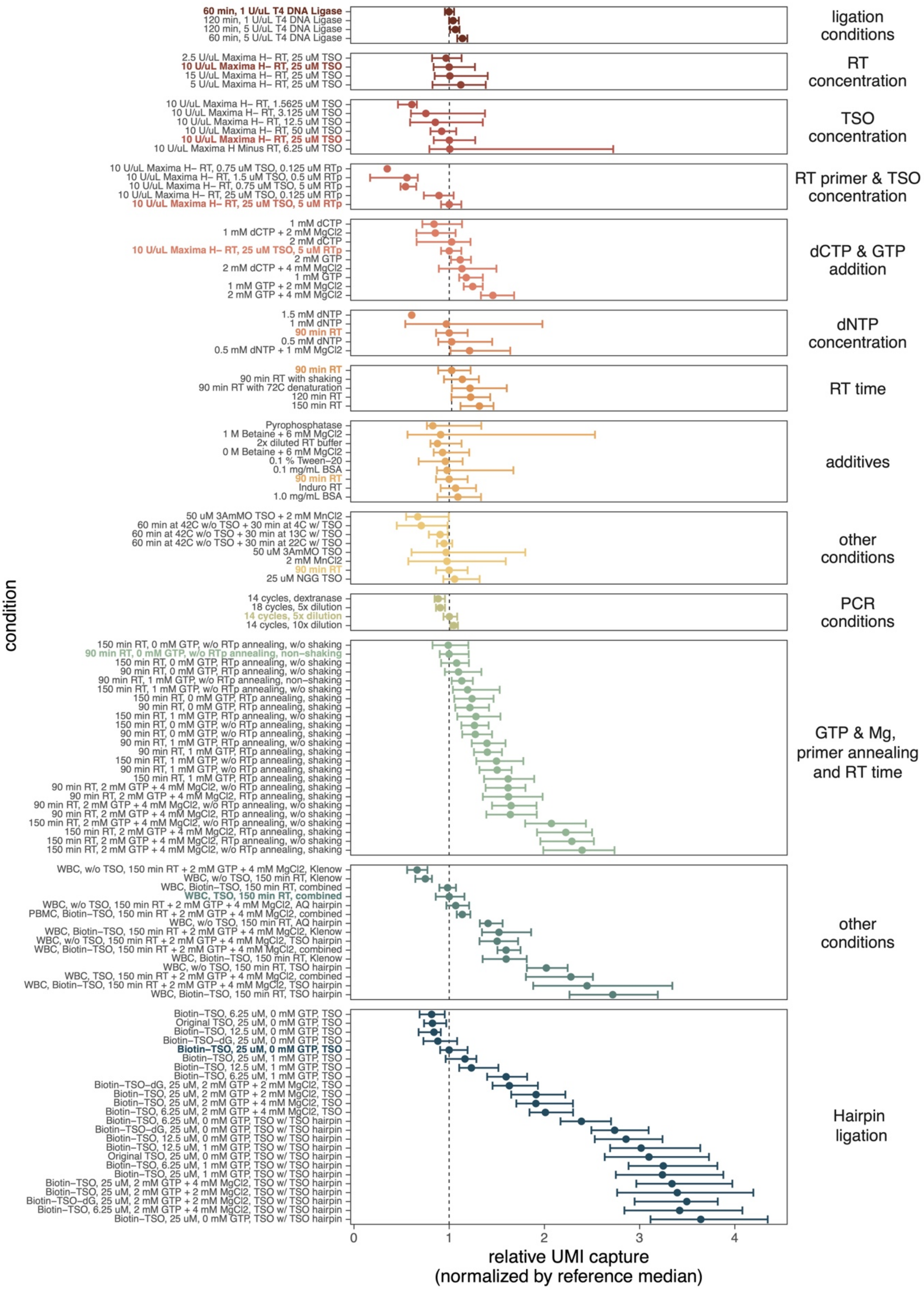


**Supplementary Figure S6. CapSeq optimization to improve UMI capture.** Each row represents a tested reaction condition. The median UMI counts at 80% sequencing saturation were calculated with 95% confidence intervals from bootstrapped distributions and normalized within condition groups (to the reference condition in bold).

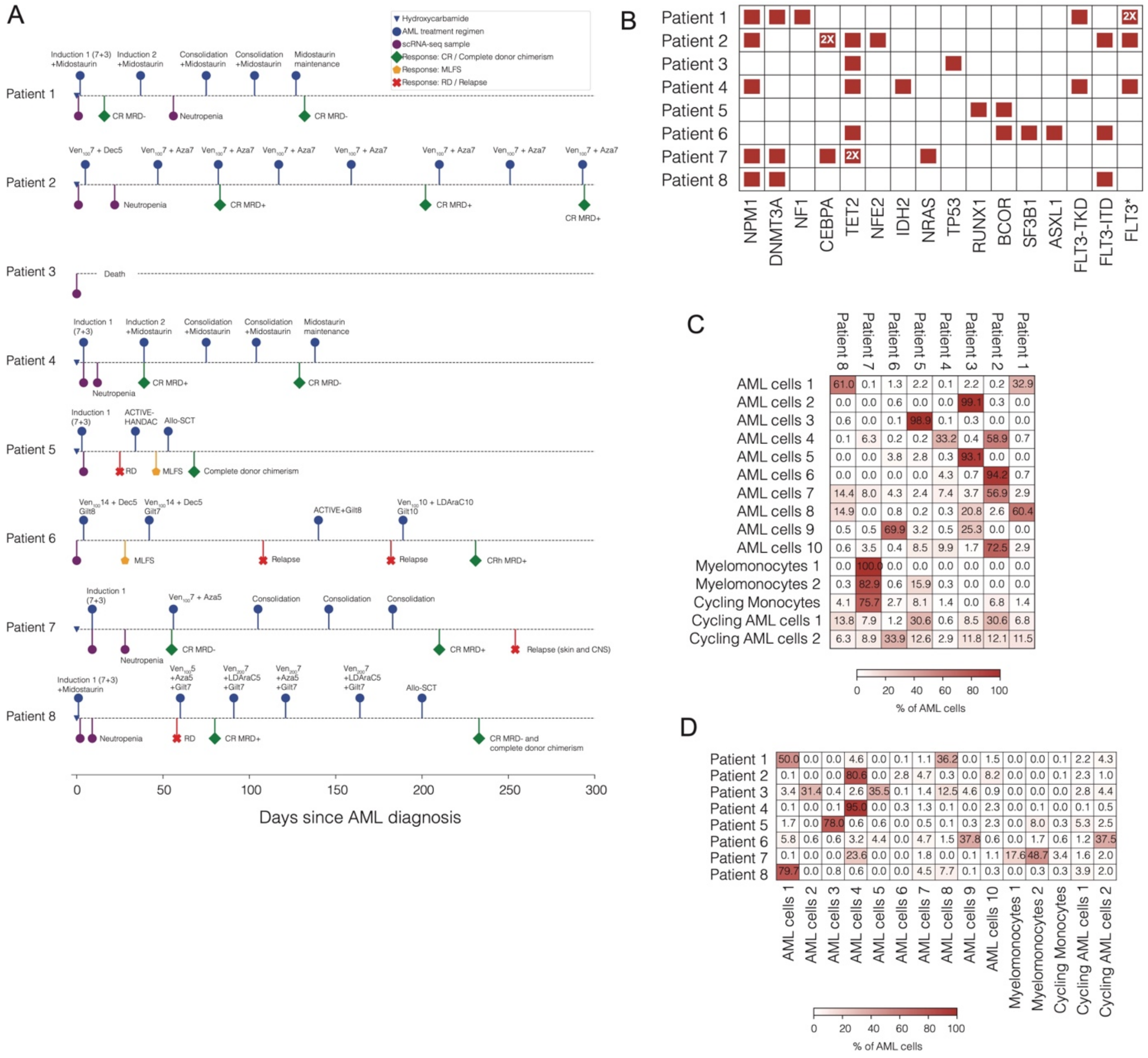


**Supplementary Figure S7. Clinical timelines of AML patients, treatment regimens, and responses over time.** **(A)** The timeline for each patient tracks their treatment history, key clinical events, and scRNA-seq sample collection points. Clinical characteristics of AML patients and details of chemotherapy regimens are provided in Table S1 and Table S2. **(B)** Mutation profile of AML patients involved in the study. **(C)** A distribution (%) of a given AML cell type in circulation across all patients. **(C)** The circulating leukemic compartment composition for each patient.

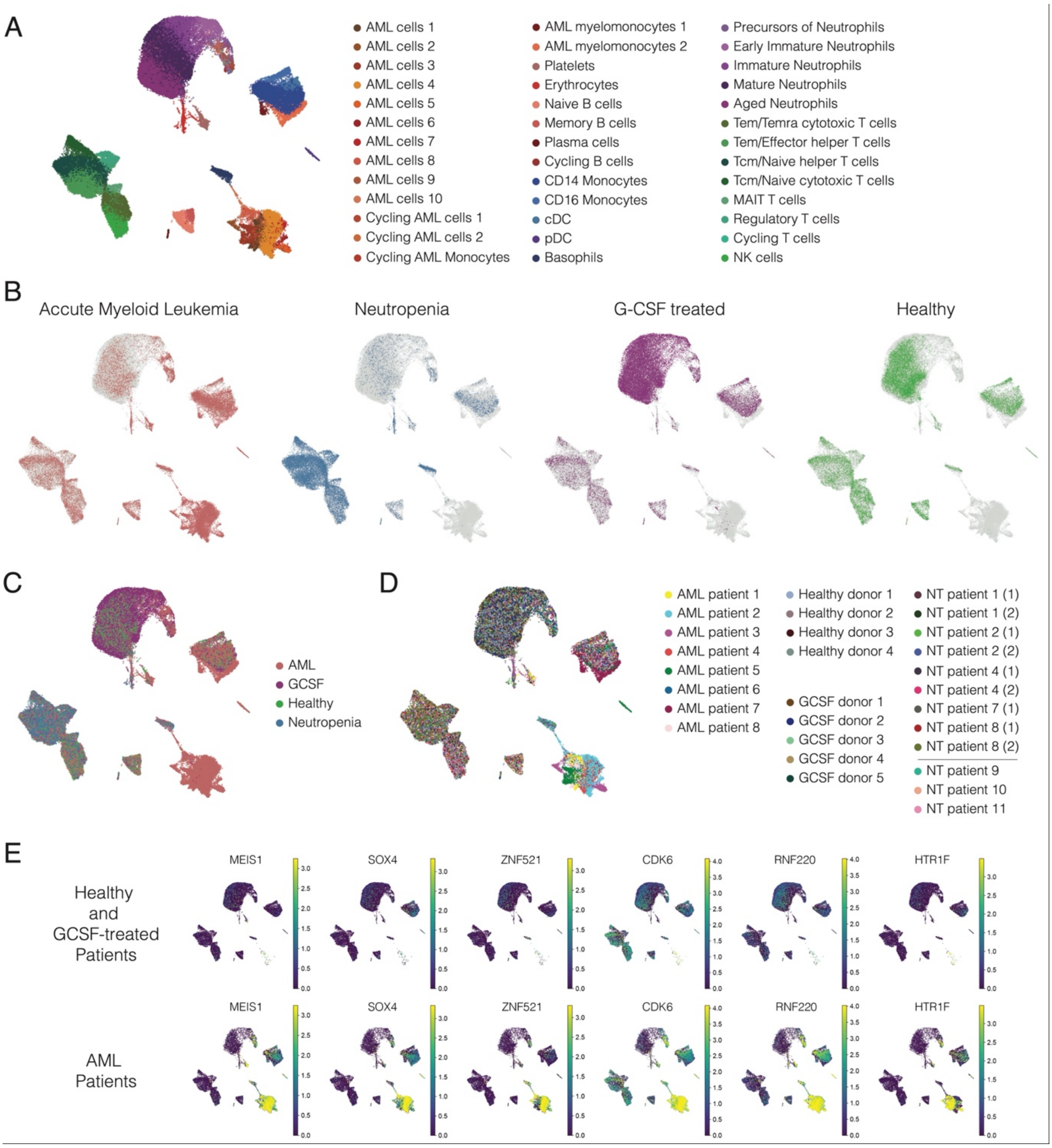


**Supplementary Figure S8. Single-cell RNA sequencing of AML, Neutropenia, G-CSF-treated, and healthy donors. (A)** UMAP visualization of cell types across all cohorts (global UMAP), showing diverse immune and AML-related cell clusters. Each color represents a different cell type. **(B)** Comparison of cell populations across different cohorts. Each panel highlights the presence and distribution of cells from each cohort on a global UMAP. **(C)** A unified UMAP plot showing cells from AML, G-CSF-treated, neutropenia, and healthy donors. Each color represents a different cohort. **(D)** Patient-specific UMAP visualization, showing the distribution of cells from AML patients, healthy donors, G-CSF-treated individuals, and neutropenia patients. Brackets indicate repeated samples from the same individual. **(E)** UMAP plots showing the log-normalized counts of selected genes (*MEIS1*, *SOX4*, *ZNF521*, *CDK6*, *RNF220*, *HTR1F*) in AML versus Healthy and G-CSF-treated individuals. Color intensity corresponds to gene expression, with scales representing values from the 0th to 99th percentile.

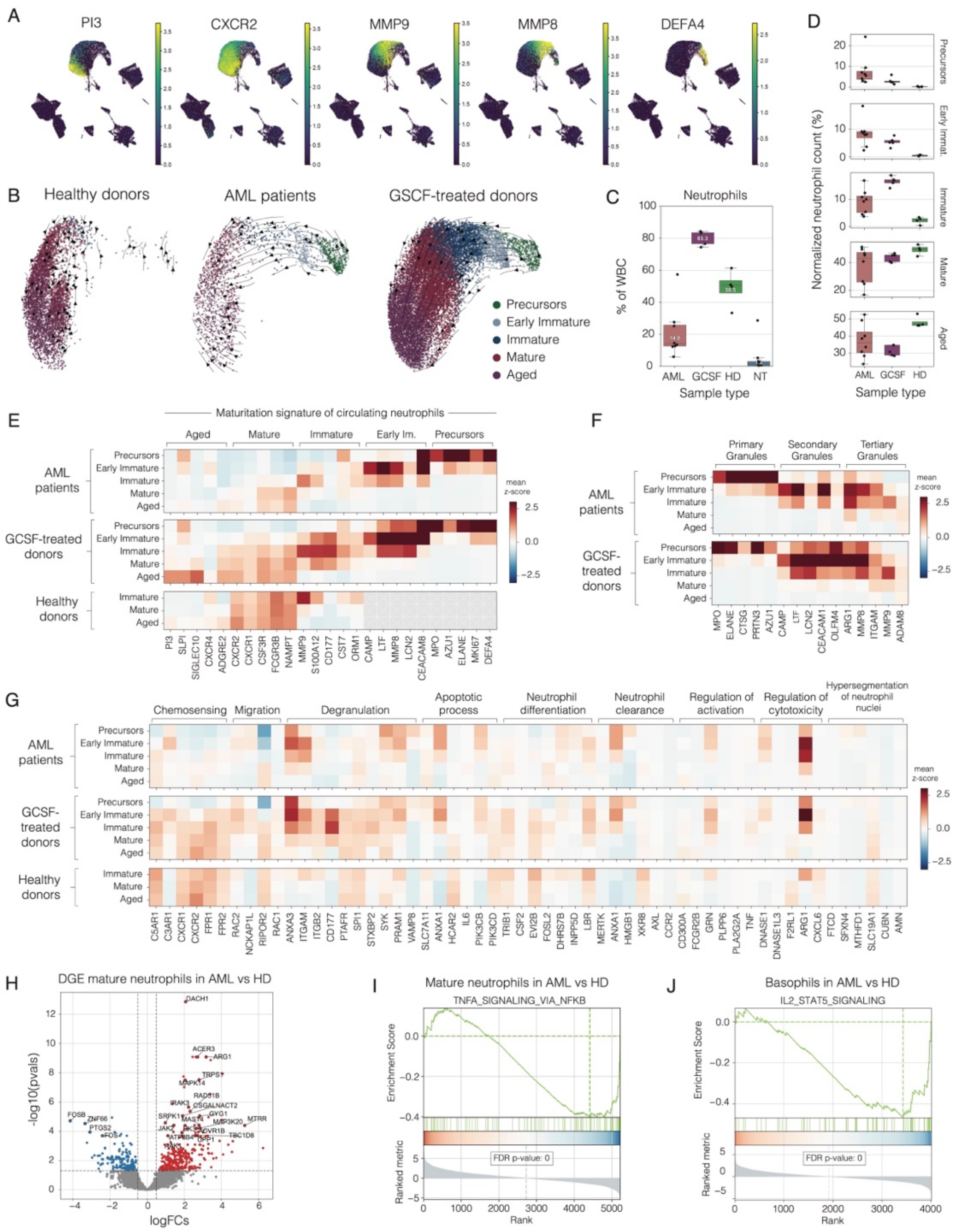


**Supplementary Figure S9. Single-cell transcriptomic profiling of neutrophil maturation across AML patients, G-CSF-treated donors, and healthy individuals. (A)** UMAP visualization of the log-normalized counts for selected gene markers (*PI3*, *CXCR2*, *MMP9*, *MMP8*, *DEFA4*) reflecting the continuum of neutrophil maturation. Color intensity corresponds to gene expression, with scales representing values from the 0th to 99th percentile. **(B)** Neutrophil RNA velocity analysis. UMAP visualization of neutrophil maturation trajectories in healthy donors, AML patients, and G-CSF-treated individuals. Arrows on the UMAPs represent RNA velocity vectors, showing the predicted direction and magnitude of transcriptional state changes. **(C)** Box plots showing neutrophil percentages within white blood cells (excluding leukemia cells) across AML, G-CSF-treated, healthy donors, and neutropenia cohorts. **(D)** Box plots showing the normalized neutrophil counts in AML, G-CSF-treated, and healthy donors across different neutrophil maturation stages (Precursors of Neutrophils, Early Immature Neutrophils, Immature Neutrophils, Mature Neutrophils, and Aged Neutrophils). **(E)** Heatmap of neutrophil maturation signature genes, showing the average expression trends across maturation stages in AML patients, G-CSF-treated individuals, and healthy donors. Color scale represents mean scaled (z-score) expression per gene. **(F)** Heatmap of primary, secondary, and tertiary granules signature genes, showing the average expression trends across neutrophil maturation stages in AML patients, G-CSF-treated individuals, and healthy donors. Color scale represents mean scaled (z-score) expression per gene. **(G)** Heatmap of the average expression of key genes involved in neutrophil related biological processes such as chemosensing, migration, degranulation, apoptosis, neutrophil differentiation, neutrophil clearance, activation regulation, cytotoxicity regulation, and nuclear hypersegmentation across AML patients, G-CSF-treated individuals, and healthy donors. Color scale represents mean scaled (z-score) expression per gene.

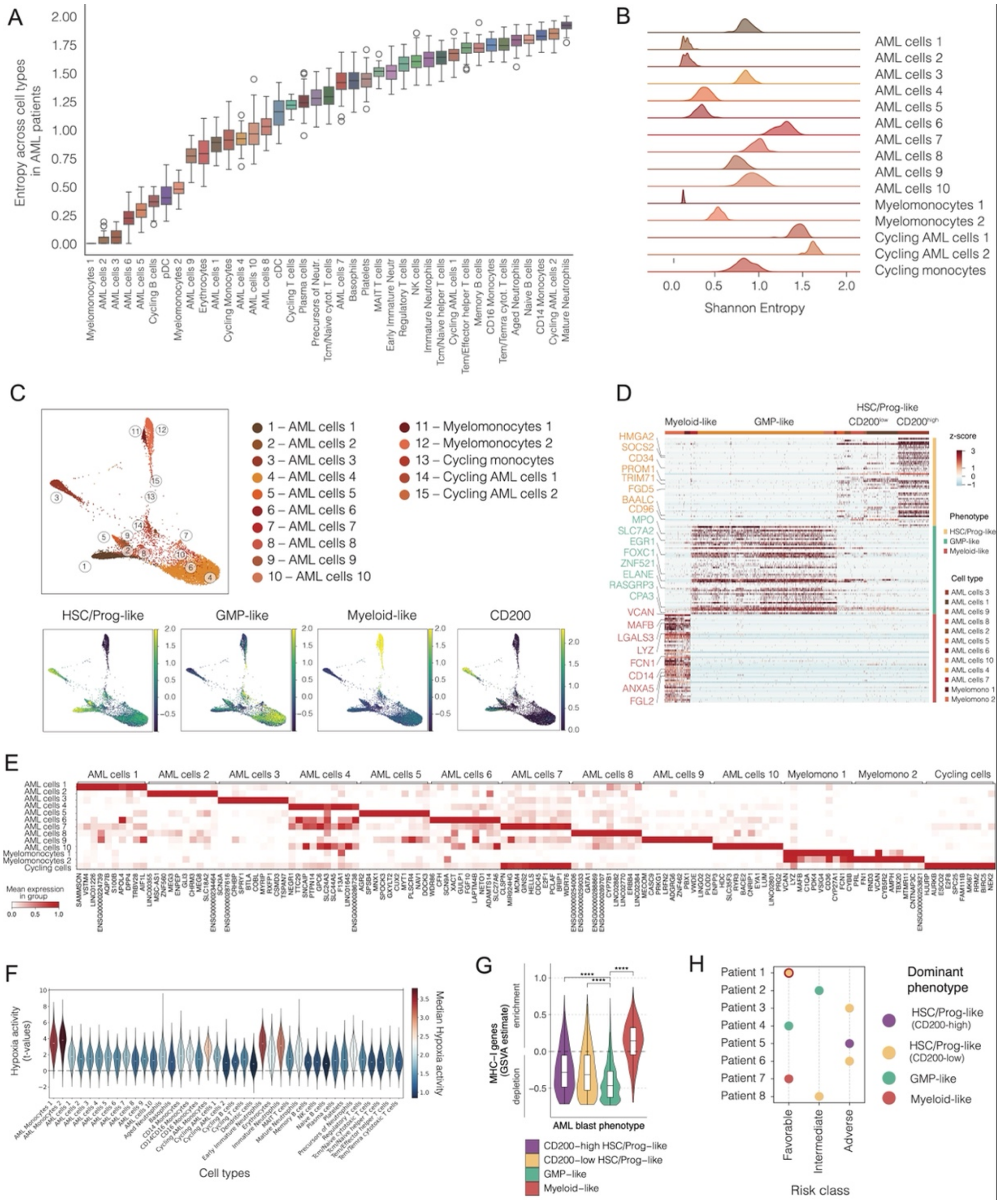


**Supplementary Figure S10. Phenotypic diversity and classification of AML blasts. (A)** Box plots showing Shannon entropy across cell types in AML patients. Lower entropy values indicate that cell phenotype is rarely shared between samples. **(B)** Density distribution of Shannon entropy in AML blasts. **(C)** Force-directed atlas (FA) visualization of AML blasts, showing average expression based on established gene signatures (see Methods) of distinct AML blasts phenotypes (HSC/Progenitor-like, GMP-like, and Myeloid-like cells), along with CD200 expression. **(D)** Heatmap of top 100 differentially expressed genes for each AML phenotype (AML phenotype vs rest of AML blasts), sorted by log2FC. DEGs were defined as those with an adjusted p-value < 0.05 and a log2 fold change (log2FC) > 1. Selected key genes (relevant in hematopoietic development) are highlighted. Color scale represents scaled (z-score) expression in cells per gene. **(E)** Heatmap of the average expression of top 10 differentially expressed genes for each AML blast type (AML blast type vs rest of AML blasts), sorted by log2FC. DEGs were defined as those with an adjusted p-value < 0.05 and a log2 fold change (log2FC) > 1. Color scale represents normalized mean expression per gene. **(F)** Violin plots of Hypoxia activity per cell type. The activity (t-values from multivariate linear model) in each cell was calculated using PROGENy approach from decoupler-py. **(G)** Depletion of MHC class I genes is pronounced in HSC/Prog-like and GMP-like cells but not in myeloid-like cells. **(H)** Association between dominant AML phenotype and ELN 2022 risk classification. Dot plot showing dominant AML phenotypes in each patient, stratified by favorable, intermediate, and adverse risk classes. Notably, Patient 1 was classified as favorable based on ELN 2022 criteria (*NPM1* mutation without *FLT3-ITD*), however, somatic *NF1* mutations have been described in literature as a factor for adverse prognosis (Eisfeld, AK., et al. Leukemia 32, 2536–2545 (2018); Tatarian et al., Blood (2022) 140 (Supplement 1): 3448–3449).

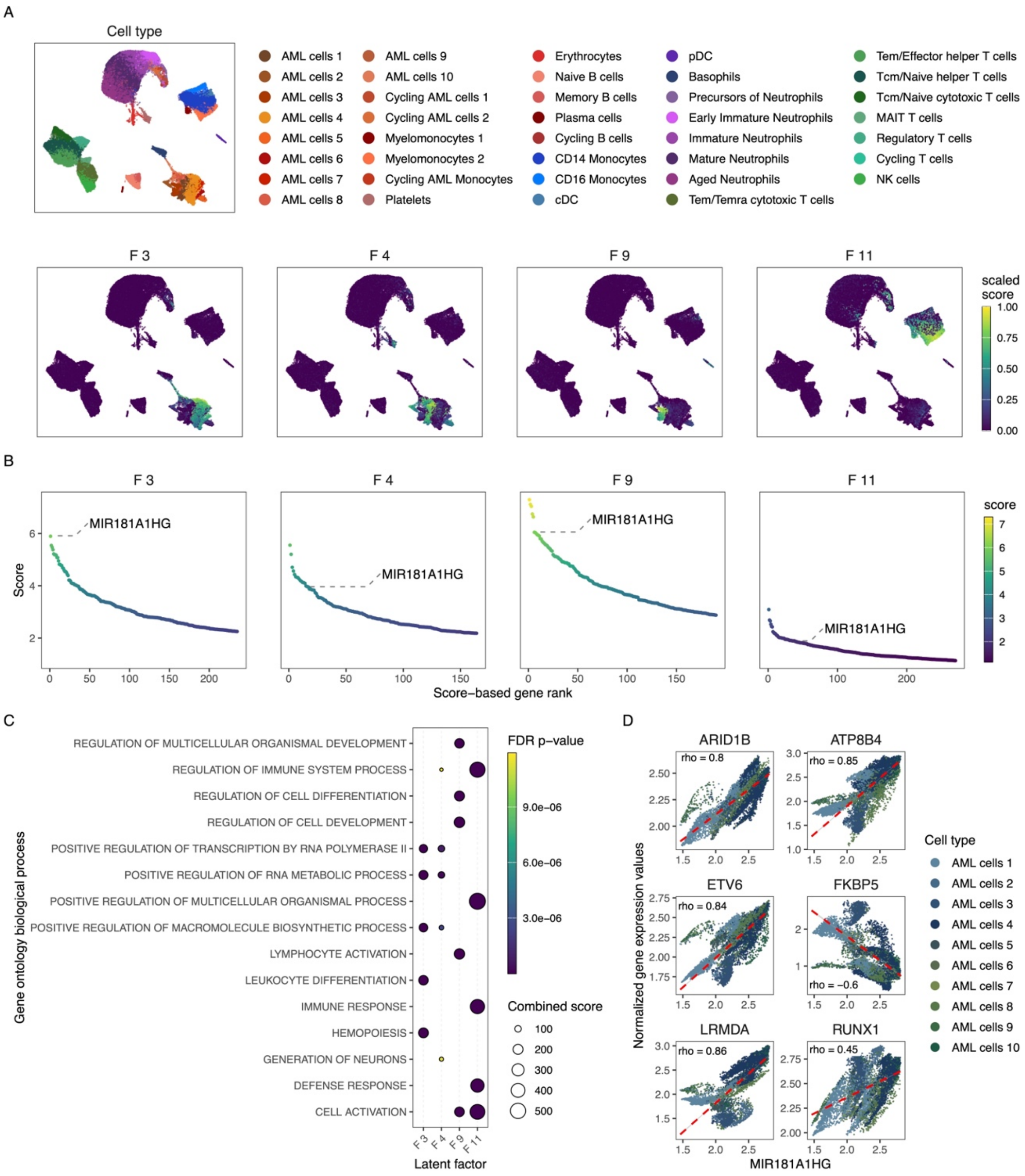


**Supplementary Figure S11. Long non-coding RNA MIR181A1HG is one of the major contributors to AML cell type-specific gene expression programs associated with cell differentiation, development, and activation. (A)** Gene expression programs (components), where *MIR181A1HG* is among the most contributing genes, are predominantly enriched in AML-related cell types. For example, F3 is enriched in AML cells 4, 6, and 7; F4 in AML cells 1, 2, and 5; F9 in AML cell 3; and F11 in AML Monocytes 1 and 2. The graph displays the scaled scores of these components in individual cells on a UMAP representation. **(B)** Genes are ranked according to their contribution to each component, with the position of the *MIR181A1HG* gene marked by a dashed line. In several components, *MIR181A1HG* ranks among the top genes. For instance, in the F4 component, it is the most contributing gene. **(C)** Functional annotation of the gene expression programs was performed using overrepresentation analysis (ORA). The analysis was performed using a list of the most contributing genes within components (gene expression programs) that include the *MIR181A1HG* gene. FDR-adjusted p-values are represented by a color gradient, while combined scores (odds ratio × -log(p-value)) are depicted proportionally by dot size. The graph displays only the top five most significant terms (based on FDR-adjusted p-values) for each component. **(D)** Exemplary cases of co-expression between MIR181A1HG and other top-ranked genes within the same components are shown. Spearman's coefficient (rho) is displayed above the linear regression curve (red dashed line). Each dot represents expression levels of the displayed genes within individual AML cells, normalized and then imputed via the MAGIC algorithm (van Dijk, David et al. Cell, V174, 716 - 729.e27).

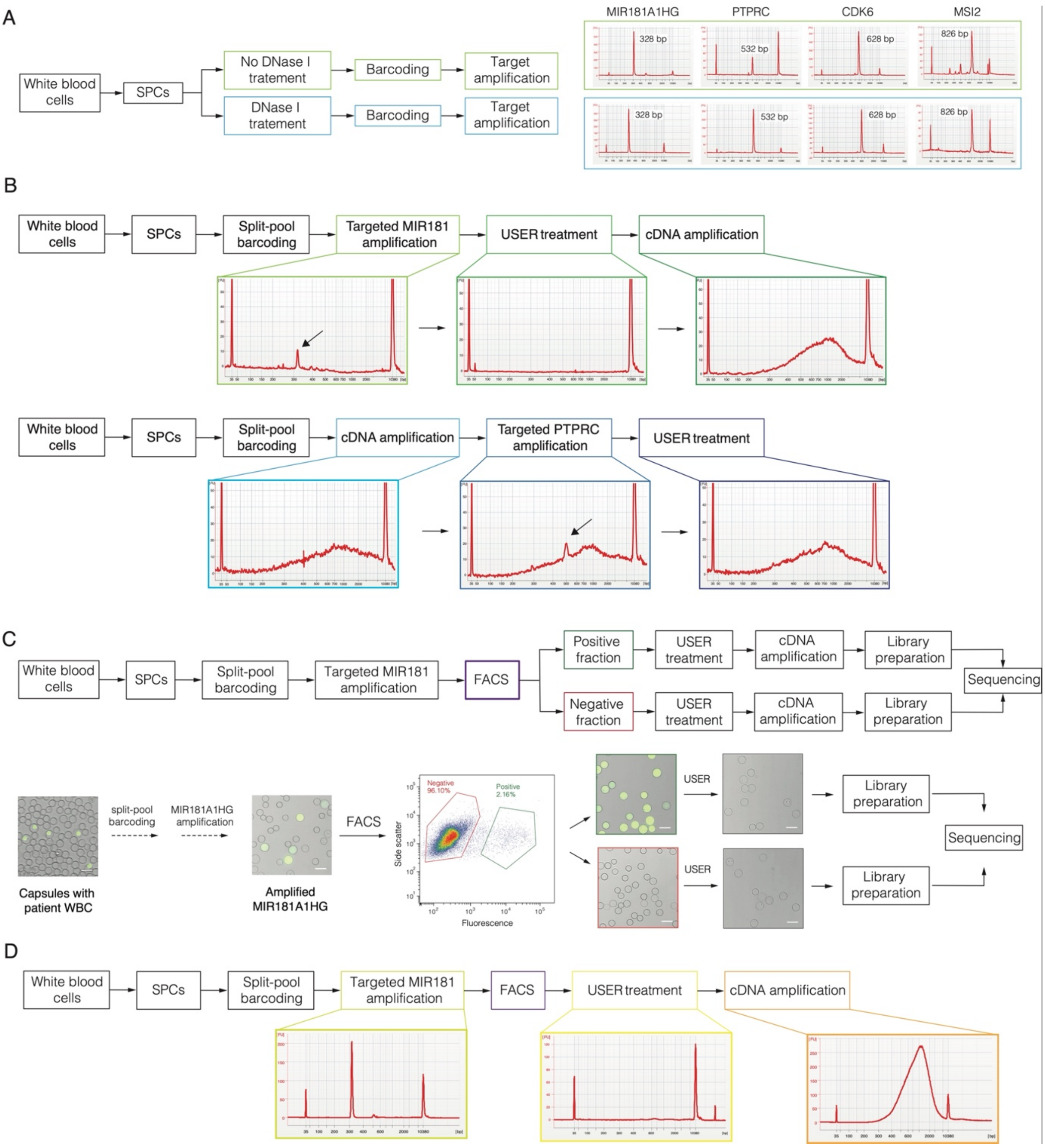


**Supplementary Figure S12. Experimental workflows for single-cell RNA cytometry and sorting based on targeted RNA expression. (A)** Comparison of DNase I treatment in target amplification. Various RNA targets (*MIR181A1HG*, *PTPRC*, *CDK6*, *MSI2*) were amplified from barcoded cDNA of white blood cells in capsules. Capsules were treated with or without DNase I prior to barcoding to validate the specificity of RT-PCR and ensure that no amplification from the genomic DNA occurs. Electropherograms show fragment sizes of amplified targets. **(B)** Target amplification and cDNA library preparation. cDNA libraries were prepared using two distinct enzymatic approaches, by amplifying either *MIR181A1HG* or *PTPRC* target. In the first approach (top), the target was amplified prior to cDNA amplification, whereas in the second approach (bottom), the target amplification was performed on already pre-amplified cDNA. The DNA trace colored by the light-blue outline contained a non-related electrical peak burst at 400 bp, which was adjusted to be smaller. Black arrows indicate the PCR amplicon **(C)** The experimental design of clinical sample sorting based on the lncRNA marker of interest *MIR181A1HG*. Sequencing libraries were prepared from both positively and negatively sorted fractions. To remove fluorescent *MIR181A1HG* amplicon, USER treatment was applied before library preparation. **(D)** Electropherograms of DNA traces at various stages of the clinical sample sorting experiment. The target lncRNA *MIR181A1HG* was amplified on barcoded, non-amplified cDNA. After sorting, fluorescent nucleic acids of the target were removed from the capsules, followed by cDNA amplification.

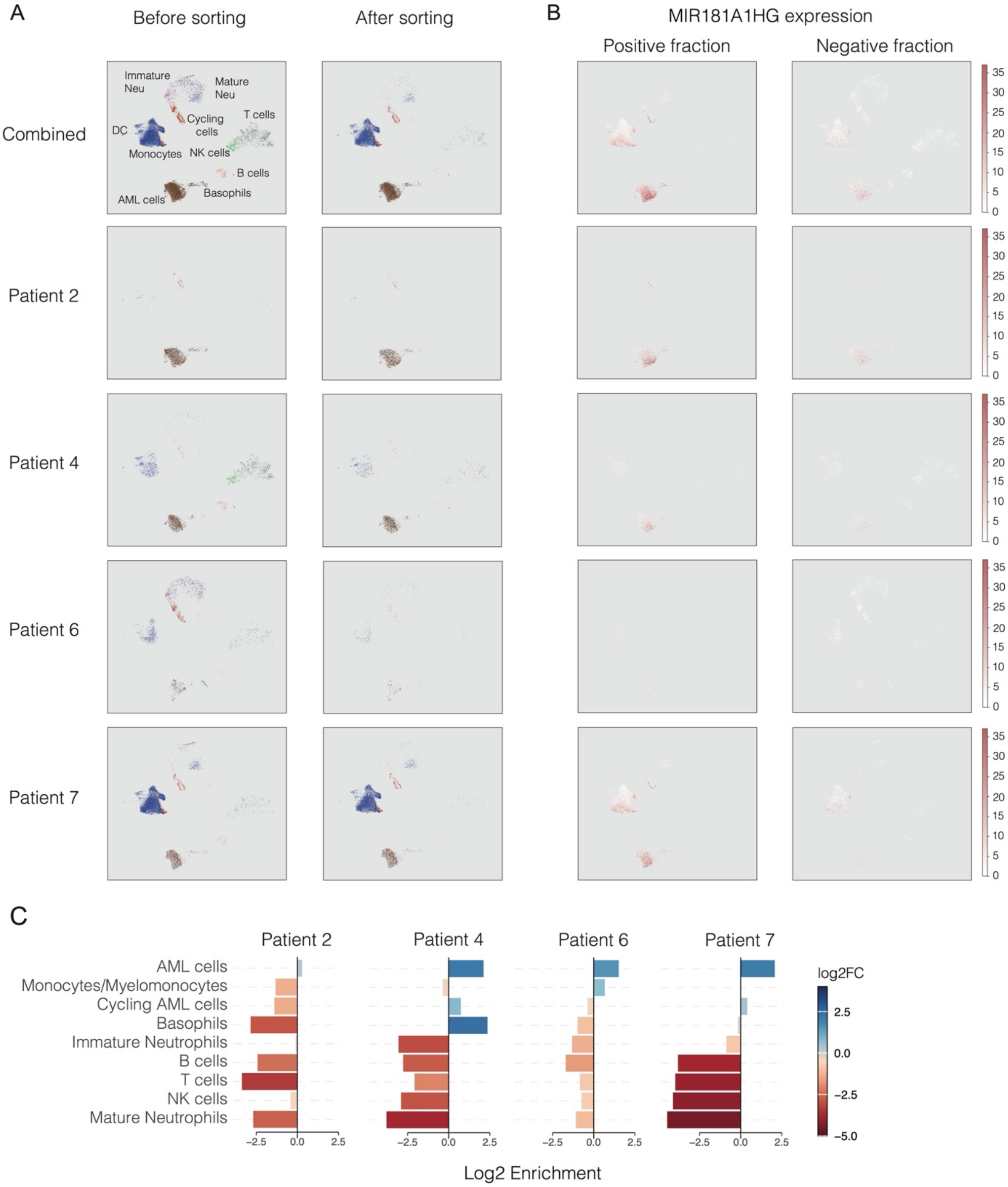


**Supplementary Figure S13. Cell type distribution and MIR181A1HG expression before and after capsules sorting on global UMAP coordinates. (A)** UMAP visualization of cell types before and after sorting. The left panel shows unsorted cells (positive and negative fractions combined), while the right panel shows the remaining cells after sorting. UMAPs are shown for the combined dataset and individual patients. **(B)** MIR181A1HG expression in sorted cell fractions. UMAP visualization of raw MIR181A1HG counts in the positive (left) and negative (right) fractions for each patient and combined dataset. **(C)** Enrichment ratio of cell types after sorting. Bar plots show the enrichment or depletion of cell types for each patient. Positive values indicate enrichment, while negative values indicate depletion.
